# Integration of Avidity and Differentiation is enabled by CD8^+^ T-cell sensing of IFN-γ

**DOI:** 10.1101/2023.03.06.531375

**Authors:** Lion F.K. Uhl, Han Cai, Jagdish N. Mahale, Andrew J. MacLean, Julie M. Mazet, Alexander J. He, Doreen Lau, Tim Elliott, Audrey Gerard

## Abstract

The most effective responses to intracellular pathogens have a breadth of T-cell clones with different affinities for their cognate peptide, and a diversity of functional phenotypes, from effector to long-lived memory cells. While high- and low-affinity T-cells are inherently skewed towards becoming effector and memory, respectively, overall, both functional subsets exploit a wide range of affinities. How the breadth of affinities and functionalities are coordinated is therefore unclear. In this study, we provide evidence that direct sensing of the cytokine IFN-γ by CD8^+^ T-cells is a factor controlling the integration of T-cell affinity and differentiation during infection. IFN-γ increases the expansion of low-affinity T-cells, allowing them to overcome the selective advantage of high-affinity T-cells. Concomitantly, IFN-γ reinforces high-affinity T-cell entry into the memory pool. As a result, direct IFN-γ sensing by CD8^+^ T-cells increases the avidity of the memory response. This comes at the expense of the primary T-cell response, for which IFN-γ decreases the avidity, leading to sub-optimum immunity to infection. IFN-γ sensing by CD8^+^ T-cells is paracrine, provided by a distinct subset of CD8^+^ T-cells called Virtual Memory T-cells, an antigen inexperienced subset that harbors memory features. Overall, we propose that IFN-γ and Virtual Memory T-cells fulfil a critical immunoregulatory role by enabling the coordination of T-cell avidity and fate.

## Introduction

CD8^+^ T-cells are critical for clearing intracellular pathogens and tumors. The efficacy of CD8^+^ T-cell responses relies on a breadth of different clones, the formation of a strong effector cytotoxic response, as well as the generation of memory cells that will provide enhanced protection upon rechallenge (Stemberger et al. 2007; Wong and Pamer 2003). T-cell differentiation into effector or memory is regulated by multiple factors, including T-cell receptor (TCR) affinity for its cognate peptide, costimulatory signals and cytokines (Kaech and Cui 2012). Despite considerable work, how those different factors are integrated to generate a robust albeit diverse short- and long-term response remains mostly unknown.

Avidity represents the capacity of T-cells to recognize infected cells and in turn elicit effector functions (Laugel et al. 2007). To estimate the avidity of an endogenous T-cell response, pMHC tetramers, which bind multivalently and often predict functional responses, are often used (Pace et al. 2012; van Gisbergen et al. 2011). But because the tetramer itself and the staining method can impact the range of discrimination between T-cell clones of different avidities (Dolton et al. 2015), this is not always used as a quantitative measure. The average avidity of the T-cell response is typically assessed by quantifying the up-regulation of activation markers (such as CD69) or effector functions (such as IFN-γ production) following stimulation of T-cells with increasing concentrations of cognate peptide (Ioannidou et al. 2017; Viganò et al. 2012). Small variations of those avidity measures in the T-cell response towards a given pathogen can have drastic consequences. For example, during COVID-19 infection, hospitalized patients have only 3-4-fold lower average T-cell avidity to spike antigens than patients with milder disease (Bacher et al. 2020). Small differences in TCR avidity can also impact tolerance, as an increase in TCR avidity of <2-fold towards a self-antigen is enough to break tolerance in a mouse model of diabetes (King et al. 2012). This indicates that the average avidity of an endogenous response has to be tightly regulated.

Together with co-receptors and co-stimulatory molecules, TCR affinity is an important factor controlling the avidity of the T-cell response (Pettmann et al. 2021). High-affinity T-cell clones are sufficient to control infections (Alexander-Miller 2005). They have a competitive advantage in interacting with antigen-presenting cells and expand better (Butz and Bevan 1998; Smith, Wikstrom, and Fazekas de 2000). Despite these selective advantages, large clonal breadth with the presence of low-affinity T-cells are consistently observed (Martinez et al. 2016; Martinez and Evavold 2015; Zehn, Lee, and Bevan 2009) during an immune response, suggesting the existence of active regulatory mechanisms.

TCR affinity regulates CD8^+^ T-cell recruitment to an immune response, but also their fate. When a single CD8^+^ T-cell clone recognizes its cognate with high-affinity, its fate is biased towards effector differentiation, whereas priming of the same clone with low-affinity peptide results in memory skewing (Kaech and Cui 2012; Solouki et al. 2020). However, a breadth of affinities is found throughout immune responses (Martinez et al. 2016; Martinez and Evavold 2015), suggesting that clonal repartition over differentiation states is actively regulated. Cells bearing the same TCR can lead to heterogenous differentiation patterns (Buchholz et al. 2013; Gerlach et al. 2013; Stemberger et al. 2007), further supporting the notion that additional factors regulate the relationship between differentiation and TCR affinity.

This overall suggests that regulation of T-cell avidity throughout an immune response requires the co-regulation of differentiation and affinity. A co-regulation implies that T-cells have to directly receive and integrate the multiple cues. IFN-γ is a cytokine that controls CD8^+^ T-cell differentiation, expansion (Gérard et al. 2013; Krummel et al. 2018; Sercan et al. 2010; Whitmire 2011; Tewari, Nakayama, and Suresh 2007; Badovinac, Tvinnereim, and Harty 2000) and the immunodominance of the T-cell response (Badovinac, Tvinnereim, and Harty 2000). Because IFN-γ regulates T-cell responses in part through direct signaling in T-cells (Gérard et al. 2013; Krummel et al. 2018; Tewari, Nakayama, and Suresh 2007; Whitmire et al. 2007), it is a strong candidate for regulating avidity throughout an immune response.

Here, we present evidence that direct IFN-γ sensing by CD8^+^ T-cells coordinate avidity and differentiation during an immune response. By deleting the receptor of IFN-γ in CD8^+^ T-cells, we demonstrated that IFN-γ limits the expansion of high-affinity T-cells, while increasing the expansion of low-affinity T-cells and skewing their differentiation towards effector. As a result, IFN-γ lowers the avidity of the primary response, while increasing the avidity of the memory response. Importantly, the regulation of avidity by IFN-γ has profound consequences, resulting in sub-optimum primary immunity towards viral infection, a trade-off leading to improved memory responses. IFN-γ signaling in CD8^+^ T-cells occurs during priming, where it is provided by a specific CD8^+^ T-cell subset, called Virtual Memory T-cells (T_VM_). We propose that T_VM_ cells fulfil a critical immunoregulatory role in modulating collective T-cell behavior. Overall, our data demonstrate that IFN-γ coordinate T-cell breadth and fate to allow for a retention of T-cell affinities throughout the immune response and a balance between short- and long-term immune responses.

## Results

### IFN-γ sensing by CD8^+^ T-cells results in sub-optimum immunity to Influenza

Given the tight regulation of T cell responses in terms of avidity and differentiation, we hypothesized that inhibiting the factors intrinsically regulating these two features would have major consequences on CD8^+^ T cell-dependent immune responses. Because IFN-γ has been implicated in controlling the immunodominance and differentiation during infection and can act directly on T-cells, we focused on IFN-γ and first determined the consequence of blocking IFN-γ sensing by CD8^+^ T-cells for viral responses. To this aim, we deleted the IFN-γR in CD8^+^ T-cells by crossing the E8I-Cre model (Maekawa et al. 2008) to IFN-γR1^flox/flox^ mice (Lee et al. 2013) (CD8-IFN-γR^KO^). IFN-γR deletion was specific for CD8^+^ T-cells, as assessed by IFN-γR1 staining (*Fig.S1A*). This was confirmed by crossing *Cd8a*-Cre to ROSA-Tomato mice, where Tomato expression was used as a read-out of Cre expression (*Fig.S1B*). CD8-IFN-γR^KO^ or control mice (WT) mice were infected with the Influenza virus strain X31 expressing OVA (X31-OVA) and their weight was monitored over time as a general read-out of control of the infection. Overall, CD8-IFN-γR^KO^ mice lost less weight than WT mice (*Fig.1A-B*) following infection with a sub-lethal dose of X31-OVA. WT mice exhibited impaired survival when infected with a higher dose of X31-OVA, while CD8-IFN-γR^KO^ mice recovered (*Fig.1C-D*). We concluded that IFN-γ-sensing by CD8^+^ T-cells results in sub-optimum effector responses against Influenza.

**Figure 1:**
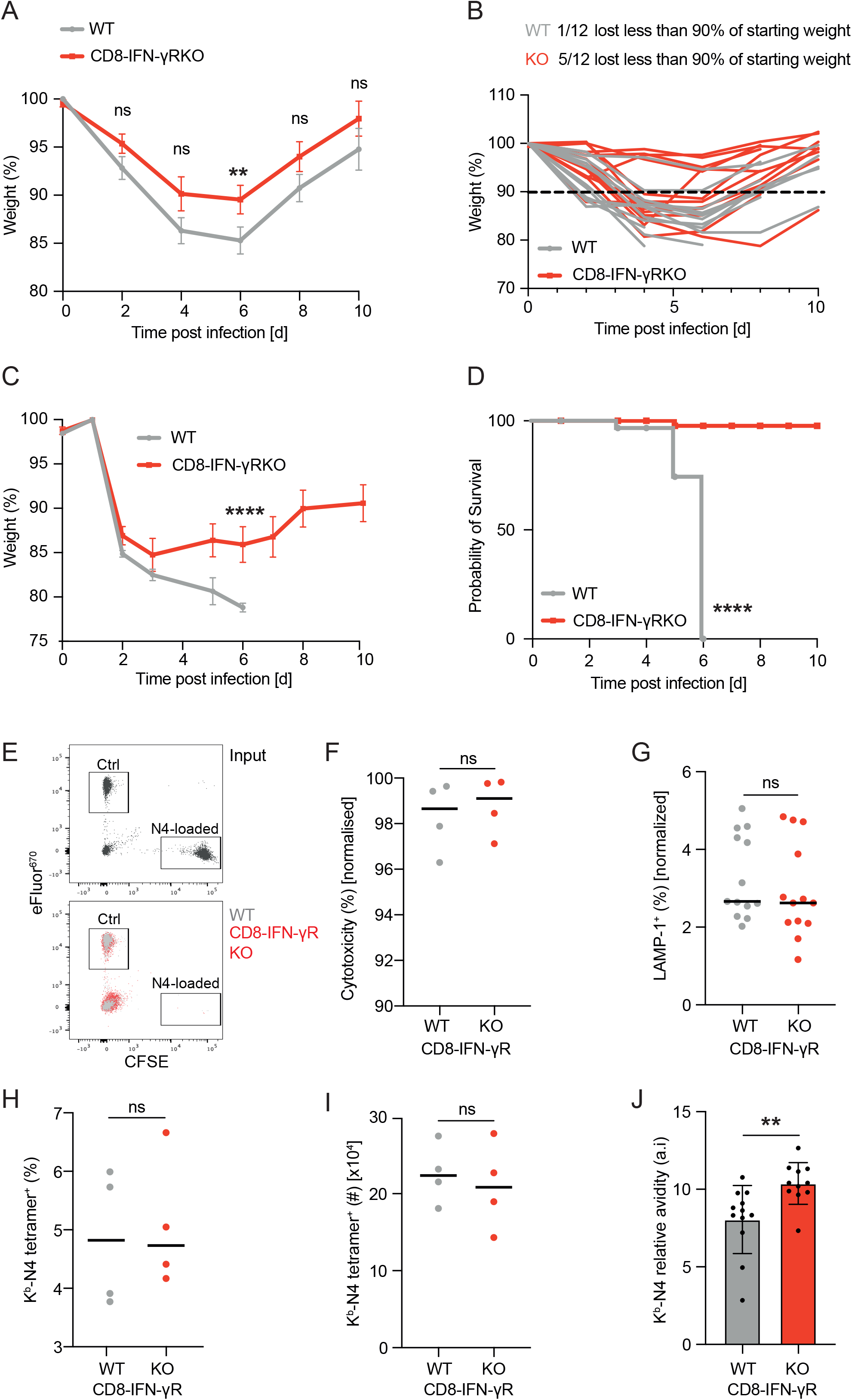
IFN-γR deletion in CD8^+^ T-cells improves effector responses against Influenza infection. **(A-B)** CD8-IFN-γR^KO^ (red) and WT (grey) mice were infected with 4×10^4^ pfu X31-OVA and weight was measured every two days to quantify relative weight loss. Graphs show average **(A)** and individual **(B)** weight loss. **(C-D)** CD8-IFN-γR^KO^ and WT mice were infected with 10^5^ pfu of X31-OVA. Graphs shows average weight **(C)** and survival **(D)** over time. **(E-F)** CD8-IFN-γR^KO^ (red) and WT (grey) mice were infected with LM-OVA and injected with differentially labelled mixtures of N4-loaded or unloaded target splenocytes at day 7 post-infection to quantify *in vivo* cytotoxicity. **E**- Dotplot example of antigen-specific in vivo cytotoxicity. **F**- Graph shows in vivo cytotoxicity normalized by the percentage of endogenous OVA-specific T-cells. **G-** CD8-IFN-γR^KO^ (red) and WT (grey) mice were infected with LM-OVA. Splenocytes were isolated between d7 and 10 and re-stimulated with N4 peptide. LAMP-1 expression in CD8^+^ T-cells was analyzed by flow cytometry normalized by the percentage of endogenous OVA-specific T-cells. **(H-J)** CD8-IFN-γR^KO^ (red) and WT (grey) mice were infected with X31-OVA and lungs were isolated 8 days post-infection to analyze the relative abundance **(H)**, absolute numbers **(I)** and relative avidity **(J)** of tetramer^+^ CD8^+^ T-cells by flow cytometry. Data are pooled from 3 (**A-D, G, J**, n = 11-13) or 4 (**F, H-I**, each dot represents an experiment) independent experiments. Error bars indicate the mean ± s.e.m **(A, C) or** median **(F-I)** or median ± s.d. (**J**). Relative avidity was calculated by dividing tetramer MFI by CD3 MFI. Cytotoxicity and LAMP-1 expression was normalized to the relative abundance of tetramer^+^ CD8^+^ T-cells within each sample. Comparison between groups was calculated using the two-tailed unpaired Student’s t-test (A-F). *P<0.05; **P<0.01; ***P<0.001; NS, not significant.

We then investigated the mechanism by which IFN-γ regulated CD8^+^ T-cell responses to Influenza infection. We tested whether increased cytotoxicity may be responsible for enhanced resistance to Influenza infection in CD8-IFN-γR^KO^ mice. Using *in vivo* cytotoxicity assay, we found that killing was efficient in both WT and CD8-IFN-γR^KO^ mice (*Fig.1E-F*). In addition, *ex vivo* surface expression of lysosome-associated membrane protein-1 (LAMP-1), a marker of cytotoxic CD8^+^ T-cell degranulation, was comparable between WT and CD8-IFN-γR^KO^ CD8^+^ T-cells (*Fig.1G*). This demonstrates that the sub-optimum CD8^+^ T-cell immunity induced by IFN-γ sensing was not the result of impaired intrinsic cytotoxic functions. Because IFN-γ is known to regulate expansion and trafficking (Bhat et al. 2017; Castro et al. 2018), we then investigated whether the increased control by CD8^+^ T cells was due to increased cell number in the lung. The number and fraction of lung OVA-specific CD8^+^ T-cells, analyzed by K^b^-N4 tetramer staining (*Fig. S1C*), at the peak of the infection was equivalent between CD8-IFN-γR^KO^ and WT T-cells (*Fig.1H-I*), indicating that IFN-γ-sensing by CD8^+^ T-cells did not regulate their overall expansion/recruitment to effector sites.

As effector function and recruitment were largely unaffected by IFN-γ sensing, we then investigated whether IFN-γ sensing by CD8^+^ T-cells affected the average avidity of the T-cell response. We analyzed K^b^-N4 tetramer binding of lung CD8^+^ T-cells and normalized the Mean Fluorescence Intensity (MFI) by CD3 expression in order to extract the relative affinity, a measure of TCR affinity and/or avidity at the population level (Pace et al. 2012). CD8-IFN-γR^KO^ T-cells exhibited increased relative affinity (*Fig.1J*), indicating that IFN-γ sensing by CD8^+^ T-cells lowers the avidity of primary responses, leading to sub-optimum immunity to Influenza.

### IFN-γ sensing by CD8^+^ T-cells decreases the avidity of the primary T-cell response

To explore whether the regulation of avidity by IFN-γ was a general mechanism, we switched to another model of infection, *Listeria monocytogenes* (LM) expressing Ovalbumin (OVA), an intracellular pathogen for which CD8^+^ T-cell response is well characterized. This model also allows us to alter the affinity and avidity of CD8^+^ T-cell responses by using LM expressing the dominant OVA peptide recognized by CD8^+^ T-cells (N4) or altered OVA peptides of lower affinity.

We first tested whether the avidity of CD8^+^ T-cell primary responses was also affected by IFN-γ sensing following LM infection. CD8-IFN-γR^KO^ or WT mice were infected with LM expressing OVA (LM-OVA) and OVA-specific CD8^+^ T-cells were analyzed by K^b^-N4 tetramer staining. In this system, expansion of endogenous OVA-specific T-cells was increased in CD8-IFN-γR^KO^ mice (*Fig.2A*), most likely because the spleen is both a priming and effector site during LM infection. In addition, OVA-specific T-cells exhibited a significant shift in tetramer binding towards higher MFI (*Fig.2B-C*). As for Influenza infection, normalizing tetramer binding by CD3 or TCR expression to extract relative avidity measures indicated that CD8-IFN-γR^KO^ CD8^+^ T-cells were of higher avidity compared to their control counterparts (*Fig.S2A-C*). This was not a consequence of increased LM-OVA load (*Fig.S2D*), increased IFN-γ production (*Fig.S2E*) or differential priming, as assessed by CD69 upregulation (*Fig.S2F*). This was also not the result of early egress of low-affinity T-cells, as the increased tetramer binding in CD8-IFN-γR^KO^ CD8^+^ T-cells was already present 5 days after infection (*Fig.S2G*).

**Figure 2:**
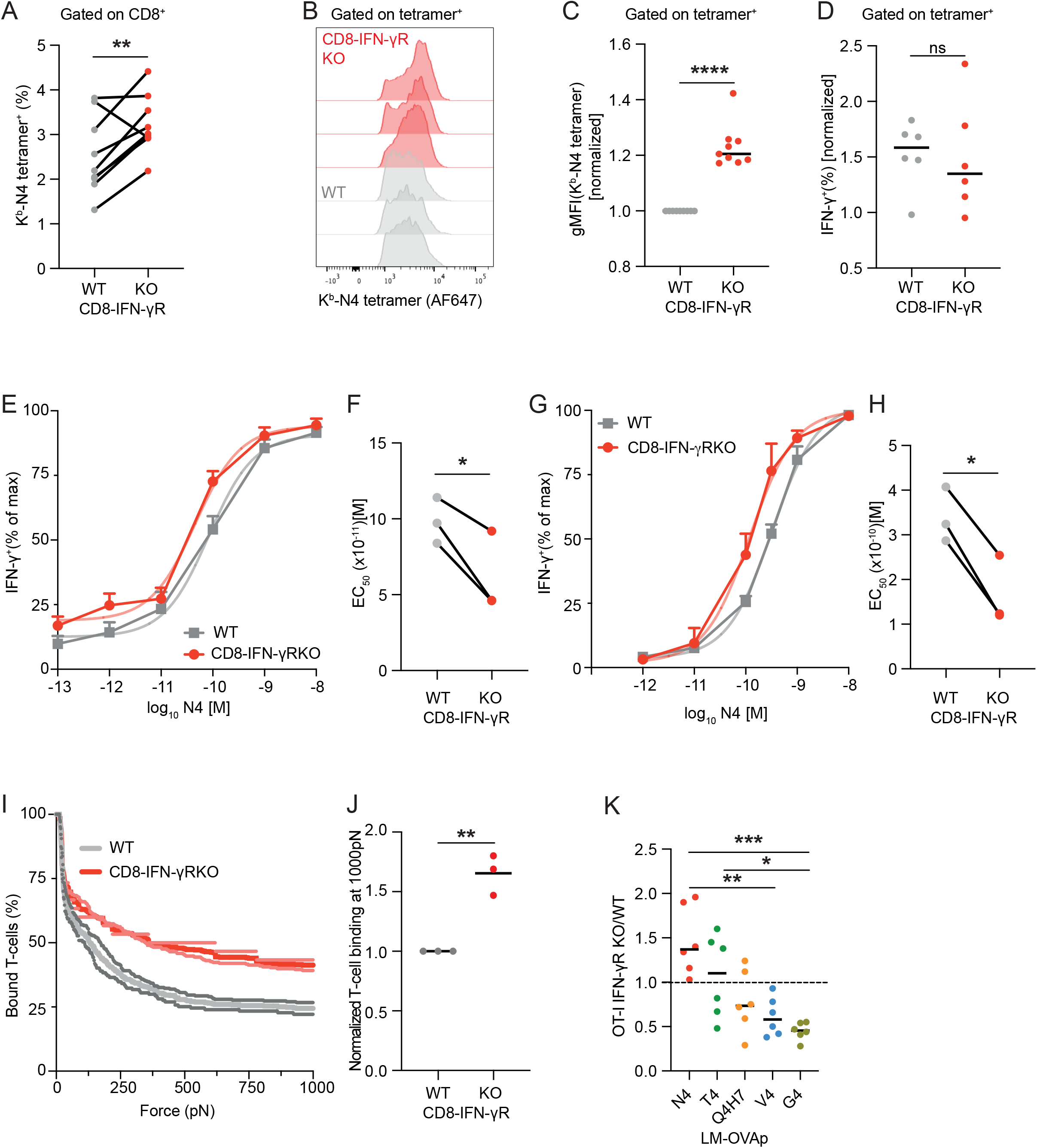
IFN-γR deletion in CD8^+^ T-cells increases the avidity of effector T-cells. **(A-H)** CD8-IFN-γR^KO^ (red) and WT (grey) mice were infected with LM-OVA and splenocytes were isolated at the peak of the response (day 7-10). (**A-C**) Endogenous OVA-specific (tetramer^+^) CD8^+^ T-cells were analyzed by flow cytometry. Graphs show their relative abundance **(A)**, representative tetramer histogram (**B**), normalized tetramer MFI (**C**). **(D-H)** Splenocytes were re-stimulated *in vitro* with the indicated concentrations of N4 for 16h. **D**-Percent of CD8-IFN-γR^KO^ (red) and WT (grey) CD8^+^ T-cells expressing IFN-γ at the maximum peptide concentration. IFN-γ expression was assessed by flow cytometry and normalized by % of tetramer^+^ cells. **(E-H)** Analysis of relative IFN-γ expression by flow cytometry **(E)** and ELISA **(G)** and quantification of EC_50_ by flow cytometry **(F)** and ELISA **(H). (I-J)** CD8-IFN-γR^KO^ and WT mice were infected with LM-OVA and tetramer^+^ CD8^+^ T-cells were sorted 9 days post-infection to measure TCR avidity after 2 days of culturing. **(I)** Analysis of CD8-IFN-γR^KO^ (red) and WT (grey) CD8^+^ T-cell binding to target cells during acoustic force measurement. **(J)** Quantification of CD8-IFN-γR^KO^ (red) and WT (grey) CD8^+^ T-cell binding at maximum force. **(K)** WT mice were transferred with 50,000 ad-mixed WT and IFN-γR^KO^ OT-I T-cells and infected with LM expressing the indicated OVA peptides. Spleens were isolated 9 days post-infection to analyze OT-I T-cells expansion by flow cytometry. Data are expressed as the ratio between IFN-γR^KO^ and WT OT-I cell number. Data are pooled from 9 (A-D, each dot represents an experiment), 3 (E-H, each dot represents an experiment), 2 (**I-J**, each dot represents an experiment) or 3 (**K**, n = 6) independent experiments. Error bars indicate the median **(C-D, J-K)** or mean ± s.e.m **(E, G)**. Tetramer-binding gMFIs of CD8-IFN-γR^KO^ samples were normalized to mean WT expression within each independent experiment. Comparison between groups was calculated using the two-tailed unpaired Student’s t-test **(A, C-D, F, H, J)** or a Two-way ANOVA and Šidák’s multiple comparison test **(K)**. *P<0.05; **P<0.01; ***P<0.001; NS, not significant.

Our data indicate that IFN-γ-sensing by CD8^+^ T-cells lowers the avidity of the primary response in two different infection models. We then sought to quantify the difference in avidity between WT and CD8-IFN-γR^KO^ CD8^+^ T-cell responses. We measured IFN-γ production following stimulation of CD8^+^ T-cells with increasing concentrations of the OVA peptide (N4) *ex vivo*. IFN-γ expression by WT and CD8-IFN-γR^KO^ T-cells were similar in magnitude (*Fig.2D*) but the EC_50_ was decreased for CD8-IFN-γR^KO^ compared to WT CD8^+^ T-cells, both by flow cytometry (*Fig.2E-F*) and ELISA (*Fig.2G-H*), indicating a higher avidity of the CD8-IFN-γR^KO^ T-cell repertoire. This demonstrated that IFN-γ-sensing by CD8^+^ T-cells decreases the avidity of the primary response by about 2-4-fold. We validated this result by quantifying OVA-specific CD8^+^ T-cell avidity by dynamic acoustic force measurements. Endogenous OVA-specific T-cells from WT and CD8-IFN-γR^KO^ mice were isolated at the peak of LM-OVA infection and allowed to adhere on a flow chamber containing a monolayer of Kb^+^, OVA-expressing cells. Increasing acoustic force was then applied and T-cell detachment from the monolayer was monitored by microscopy. Using this method, we confirmed that CD8-IFN-γR^KO^ OVA-specific T-cells displayed increased avidity compared with WT T-cells (*Fig.2I-J*). Taken together, we demonstrated that IFN-γ-sensing by CD8^+^ T-cells decreases the avidity of the primary response using multiple techniques, showing that this phenotype is robust.

To understand whether the function of IFN-γ on the avidity of the CD8^+^ T-cell response could originate from modulating the behavior of clones with different TCR affinity, we fixed the CD8^+^ T-cell clonotype by using the OVA-specific CD8^+^ T-cell clone OT-I, and varied the affinity of TCR priming. We transferred the smallest number of OT-I T-cells allowing reliable detection following priming with peptides exhibiting low-affinity for the OT-I TCR. WT and IFN-γR^KO^ OT-I were co-transferred in mice that were subsequently challenged with LM expressing either the high-affinity OVA peptide N4 or altered peptides of lower affinity. We then compared the expansion of WT and IFN-γR^KO^ OT-I T-cells upon priming. Expansion of IFN-γR^KO^ OT-I T-cells was increased following high-affinity (N4 peptide) priming at the peak of the response compared to WT. Surprisingly, priming with low-affinity peptides led to decreased frequency of IFN-γR^KO^ over WT OT-I T-cells (*Fig.2K, S2H*), demonstrating that direct sensing of IFN-γ by CD8^+^ T-cells differentially affected T-cell expansion according to TCR affinity, favoring the participation of low-affinity T-cells while restraining the expansion of high-affinity T cells. This is consistent with our finding that IFN-γ-sensing by CD8^+^ T-cells decreases the avidity of the primary response.

Altogether, we concluded that IFN-γ signaling in CD8^+^ T-cells curtails expansion of T-cells with high affinity TCR, resulting in decreased avidity of the primary response.

### CD8^+^ T-cell paracrine IFN-γ signalling during priming regulates the avidity of the primary response

IFN-γ production during LM-OVA infection is characterized by an early wave of IFN-γ production occurring during priming and a second wave during the peak of the effector response (Krummel et al. 2018). To determine when IFN-γ was sensed by CD8^+^ T-cells to regulate T-cell avidity, anti-IFN-γ was administered either 16-24h or at day 5 and 6 post-infection. Similar to deleting IFN-γR on CD8^+^ T-cells, blocking IFN-γ 16-24h post-infection resulted in increased abundance of OVA-specific CD8^+^ T-cells (*Fig.3A*) and increased tetramer (but not CD3) staining (*Fig.3B-C*). Taken together, we concluded that IFN-γ was sensed 16-24h post-infection to regulate the expansion and avidity of OVA-specific CD8^+^ T-cells. Blocking IFN-γ during the second wave had no effect on those measures.

**Figure 3:**
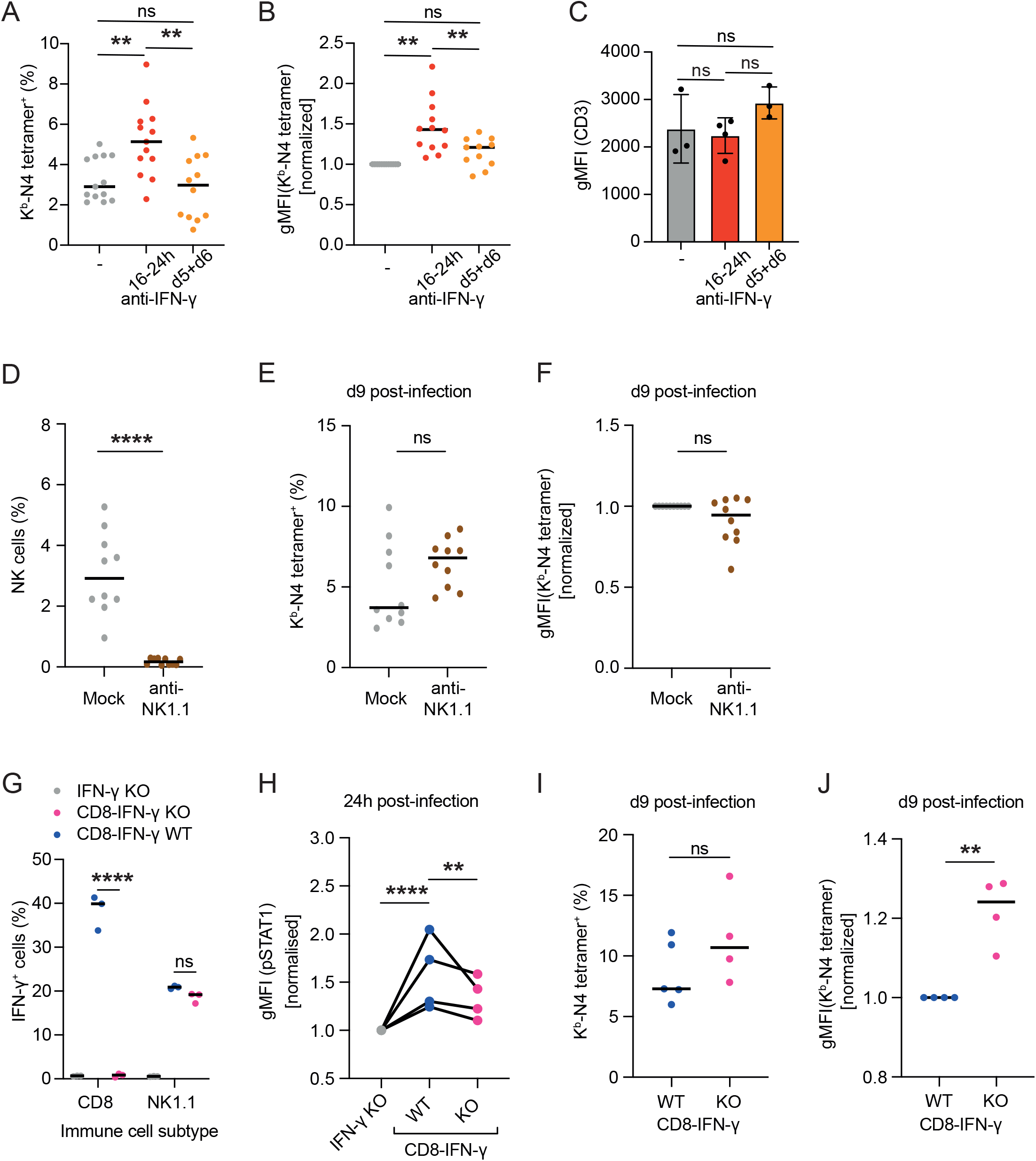
IFN-γ-is by CD8^+^ T-cells at priming and is paracrine. **(A-C)** WT mice were infected with LM-OVA and mice were either left untreated (grey) or treated with anti-IFN-γ 16-24 hours (red) or at day 5 and 6 (orange) post-infection. Splenocytes were isolated 9 days post infection and analyzed by flow cytometry. Graphs show relative abundance **(A)**, tetramer gMFIs **(B)**, and CD3 gMFI **(C)** of tetramer^+^ CD8^+^ T-cells. **(D-E)** WT mice were treated with depleting NK1.1 antibodies (brown) or control antibodies (grey) and infected with LM-OVA. **D-** Splenocytes were isolated after 24h and the proportion of NK cells was quantified by flow cytometry. **(E-F)** Splenocytes were isolated after 9 days and tetramer^+^ CD8^+^ T-cells were analyzed by flow cytometry. Graphs show relative abundance **(E)** and tetramer gMFIs **(F)** of tetramer^+^ CD8^+^ T-cells. **(G-H)** WT (blue) and CD8-IFN-γ^KO^ (pink) chimera mice were infected with LM-OVA. **G-** Splenocytes were isolated after 24h and stimulated *in vitro* with PMA/Ionomycin. IFN-γ expression by NK and CD8^+^ T-cells was quantified by flow cytometry. **H-** Graph shows the percentage of IFN-γ-expressing CD8^+^ T-cells or NK cells. Splenocytes were isolated after 24h and STAT1 phosphorylation in activated (CD69^+^) CD8^+^ T-cells was analyzed by flow cytometry. **(I-J)** Splenocytes were isolated after 9 days and tetramer^+^ CD8^+^ T-cells were analyzed by flow cytometry. Graphs show relative abundance **(I)** and tetramer gMFIs **(J)** of tetramer^+^ CD8^+^ T-cells. Data are pooled from 3 (**A-C**, n = 6-12), 2 (**D-F**, n = 10), 4 (**G-J**, each dot represents an experiment) independent experiments. STAT1 phosphorylation was normalized to control (IFN-γKO) pSTAT1 MFI. Error bars indicate the median. Comparison between groups was calculated using the two-tailed unpaired Student’s t-test **(D-F, I-J)** or a Two-way ANOVA and Šidák’s multiple comparison test **(A-C, G-H)**. *P<0.05; **P<0.01; ***P<0.001; NS, not significant.

Because NK cells were the main source of IFN-γ at the onset of LM infection (*Fig.S3*), we investigated whether they contributed to the regulation of T-cell avidity. To do so, we ablated NK cells with depleting NK1.1 antibody (*Fig.3D*) and infected mice with LM-OVA. The proportion and avidity of the T-cell response against OVA was assessed at the peak of the response. Depleting NK cells resulted in an increase in the proportion of tetramer^+^ T-cells (*Fig.3E*), most likely due to the increased bacterial load, as mice without NK cells fail to control LM infection (Viegas et al. 2013). However, this was not accompanied by an increase in tetramer staining (*Fig.3F*), demonstrating that NK cells were not the source of IFN-γ implicated in the regulation of T-cell avidity.

Because IFN-γ is shared between CD8^+^ T-cells during priming (Krummel et al. 2018) and the second cellular source of IFN-γ (*Fig.S3*), we hypothesized that CD8^+^ T-cell-derived IFN-γ may be the dominant source regulating CD8^+^ T-cell avidity. To test this, we created mixed bone marrow (BM) chimeras by reconstituting lethally irradiated IFN-γ^KO^ mice with ad-mixed IFN-γ^KO^ and CD8α^KO^ BM, resulting in IFN-γ deletion specifically in CD8^+^ T-cells (CD8-IFN-γ^KO^ mice). Reconstitution of IFN-γ^KO^ mice with ad-mixed IFN-γ^KO^ and WT BM was used to generate control (WT) mice (*Fig.3G*). STAT1 phosphorylation in CD8^+^ T-cells was analyzed 24h post-LM-OVA infection, as a read-out of IFN-γ signaling. WT CD8^+^ T-cells exhibited enhanced IFN-γ signaling compared to CD8-IFN-γ^KO^ cells (*Fig.3H*). Ablation of CD8-induced IFN-γ led to 40% inhibition of IFN-γ signaling (*Fig.3H*) while CD8^+^ T-cells constitute a small proportion of IFN-γ-producing cells (around 15%, *Fig.S3*), suggesting that CD8^+^ T-cells are highly sensitive to their own IFN-γ. At the peak of the response, we observed an increase in the proportion of OVA-specific T-cells (*Fig.3I*) in CD8-IFN-γ^KO^ mice, with enhanced tetramer binding (*Fig.3J*), as observed with CD8-IFN-γR^KO^ mice (*Fig.2C*).

Taken together, we concluded that CD8^+^ T-cell paracrine IFN-γ signaling regulates the avidity of their response.

### IFN-γ is provided by Virtual Memory T (T_VM_) cells during priming

Given that CD8^+^ T-cells primarily sense their own IFN-γ to regulate their avidity, we then characterized which CD8^+^ T-cell subset secreted IFN-γ using scRNA-seq of CD8^+^ T-cells from control mice and mice infected with LM-OVA for 24h. Unsupervised hierarchical clustering identified 6 clusters (*Fig.S4A*), which were labelled based on known markers of naïve and memory T-cells (*Fig.S4B*). We distinguished 4 clusters harboring a naïve phenotype that we merged (*Fig.4A-B*), and 2 clusters with a memory phenotype (*Cd44, Eomes* and *Cxcr3*). One memory cluster had features of naïve cells, such as *Lef1, Ccr7* and *Sell* (encoding for CD62L) expression, expressed high levels of *Il2rb* and low levels of the integrin *Itga4* (*Fig.4A-B*). These markers are characteristic of Virtual Memory T-cells (T_VM_), a subset of antigen-independent memory T-cells (Hussain and Quinn 2019). The T_VM_ characteristic expression pattern was confirmed by Differential Gene Expression analysis (*Fig.4C*). Analysis of IFN-γ transcript revealed that T_VM_ cells were the predominant subpopulation producing IFN-γ (*Fig.4D-E*). We confirmed this by flow cytometry using GREAT mice, an IFN-γ reporter strain whereby YFP expression is driven by the IFN-γ promoter (Reinhardt, Liang, and Locksley 2009). GREAT mice were infected with LM-OVA and the production of IFN-γ by the different CD8^+^ T-cell subsets was assessed using CD44, CD122 and CD49d *(Itga4)* to delineate naïve, T_VM_ and antigen-experienced memory CD8^+^ T-cells. Up to 70% of IFN-γ producers were T_VM_ (*Fig.4F*) 24h after LM-OVA infection. Similar data was observed with LM expressing gp33, ruling out the possibility of antigenic bias (*Fig.4F*). We concluded that T_VM_ was the main CD8^+^ T-cell subset producing IFN-γ during priming.

**Figure 4:**
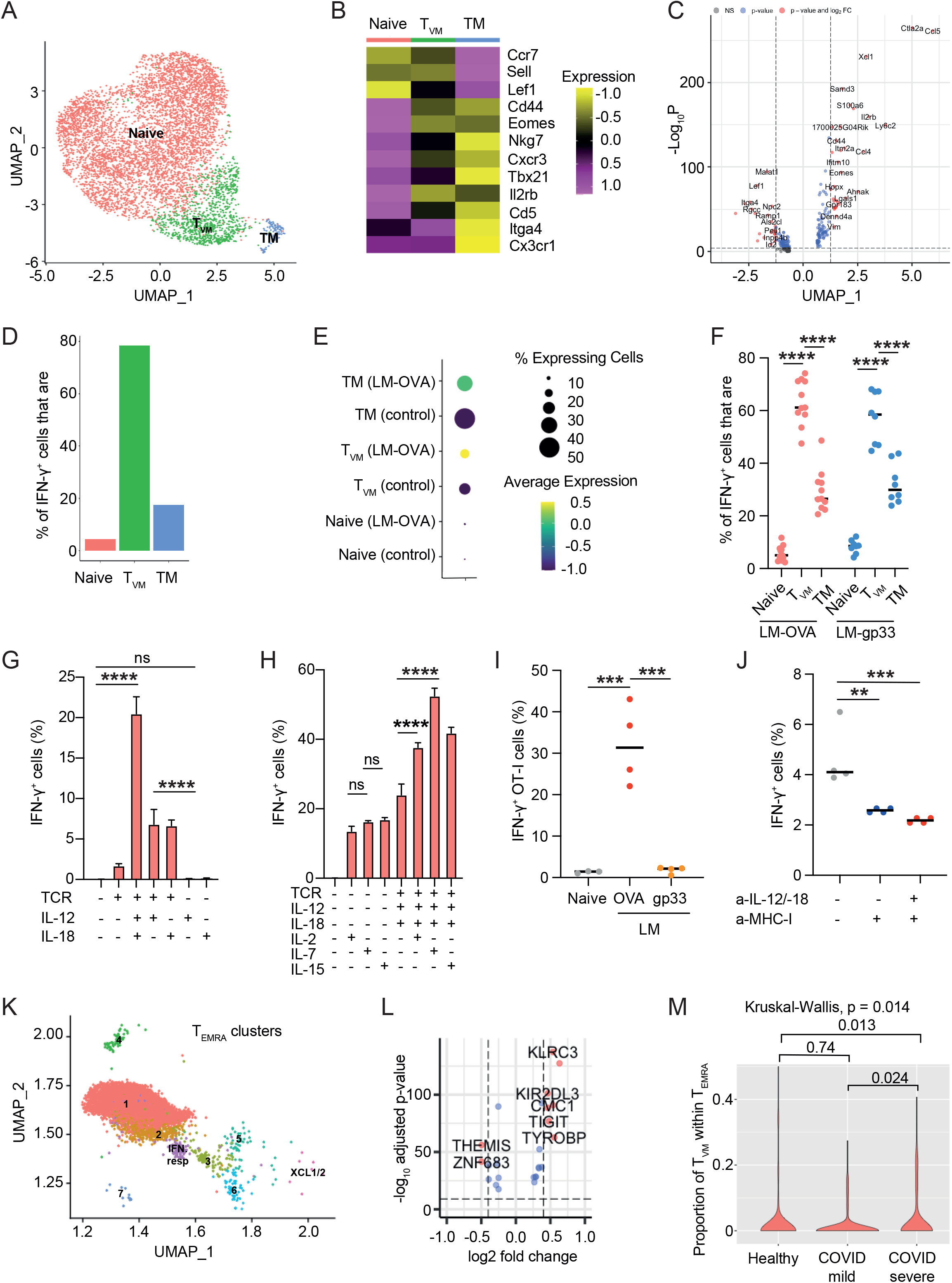
T_VM_ are the main CD8^+^ T-cell source of IFN-γ-during priming. **(A-F)** CD8^+^ T-cells from naïve mice or mice infected with LM-OVA for 24h were sorted and subjected to scRNA-seq analysis (n=3). **A-** Graph-based clustering (n=3224 ctrl; n=3495 LM-OVA) of the assembled cell states. **B-** Heatmap shows the average expression of selected markers. **C-** Volcanoplot shows specific differentially regulated genes between T_VM_ and other CD8^+^ T-cells. Green dots: genes with log2 (fold-change) value >0.5 or <-05; Blue dots: genes with an adjusted p value<0.05; Red dots: genes with log2 (fold-change) value >0.5 or <-05 and an adjusted p value<0.05. **D**-Graph shows the relative frequency of naïve, T_VM_ or memory T cells (TM) CD8^+^ T-cells within the IFN-γ^+^ CD8^+^ T-cell fraction. **E-** Dot plot shows the average expression of IFN-γ in naïve, T_VM_ or TM cells at steady-state and 24h post LM-OVA infection. **F-** GREAT mice were infected with either LM-OVA (red) or LM-gp33 (blue) and CD8^+^ T-cell populations among IFN-γ (YFP)^+^ splenocytes were analyzed by flow cytometry 24h after infection. **(G-H)** CD8^+^ T from GREAT mice were stimulated *in vitro* with anti-CD3/-28 (TCR), and the indicated cytokines. IFN-γ production was analyzed by flow cytometry after 24h. **I-** Mice were transferred with 2×10^6^ GREAT OT-I CD8^+^ T-cells and infected with LM-OVA (red) or LM-gp33 (orange). IFN-γ (YFP) expression by OT-I was quantified by flow cytometry 24 hours post-infection. **J-** GREAT mice were infected with LM-OVA and treated with anti-MHC class-I, and anti-IL-12/-18 as indicated. IFN-γ (YFP) production was quantified in CD8^+^ T-cells 24h post-infection. **(K-M)** T_EMRA_ clusters from the COMBAT human blood atlas were reanalyzed according to COVID-19 severity. **K-** Graph-based clustering of the T_EMRA_ clusters. **L-** Volcanoplot shows specific differentially regulated genes between T_VM_ and other T_EMRA_. Green dots: genes with log2 (fold-change) value >0.5 or <-05; Blue dots: genes with an adjusted p value<0.05; Red dots: genes with log2 (fold-change) value >0.5 or <-05 and an adjusted p value<0.05. **M-** Violinplot shows the percentage of T_EMRA_ that are T_VM_ according to disease state and severity. Each dot is a patient. Pairwise group comparisons with two-sided Wilcoxon signed-rank test. Data are pooled from 3 (**F-H**, n = 6-12) or 4 (**I-J**, each dot represents an experiments) independent experiments. Error bars indicate the median **(F, I-J)** or mean ± s.e.m **(G-H)**. *P<0.05; **P<0.01; ***P<0.001; NS, not significant.

T_VM_ cells are known to exhibit enhanced sensitivity to inflammatory cytokines, resulting in bystander activation (White et al. 2016). Differential pathway analysis, however, suggested that T_VM_ priming was more complex than bystander priming during LM-OVA infection. While true (antigen-experienced) memory T-cells displayed signatures of cytokine priming, T_VM_ exhibited additional signatures, some shared with naïve T-cells, including increased metabolism, heat-shock response, and cell cycle, which could indicate TCR priming (*Fig.S4C*). To address whether IFN-γ production by T_VM_ required TCR priming, CD8^+^ T-cells from GREAT mice were activated *in vitro* with anti-CD3ε/-CD28 (“TCR stimulation”) and/or inflammatory cytokines, and IFN-γ expression was assessed after 24h. While IL-12 and IL-18 treatment induced some level of IFN-γ production, adding TCR priming synergized with IL-12/18 (*Fig.4G*). IL-2, IL-7, and IL-15, which share the same γ-chain receptor, further synergized with TCR and IL-12/18 stimulation to increase IFN-γ (*Fig.4H*). Other known inducers of IFN-γ production did not lead to IFN-γ production by CD8^+^ T-cells (*Fig.S4D*). This suggests that maximum

IFN-γ production by T_VM_ required TCR priming. We validated this *in vivo*. We crossed GREAT mice with OT-I mice to generate GREAT OT-I T-cells, which were transferred in WT recipients. Mice were then infected with either LM-OVA or LM-gp33. For this, we increased the frequency of OT-I cells transferred in order to detect OT-I T_VM_. IFN-γ expression by T_VM_ OT-I T-cells was observed upon infection with LM-OVA but not LM-gp33 (*Fig.4I*), confirming the requirement of TCR priming for early IFN-γ induction *in vivo*. In addition, we confirmed that IFN-γ production by CD8^+^ T-cells required TCR priming *in vivo* by using Nur77-reporter mice, for which GFP expression is controlled by the Nur77 promoter and correlates with TCR priming (Moran et al. 2011). Mice were infected with LM-OVA and after 24h, we observed a significant shift in GFP expression in IFN-γ^+^ CD8^+^ T-cells compared to non-producers (*Fig.S4E*). Finally, blocking MHC class-I 24 hours post-infection also significantly decreased IFN-γ expression by CD8^+^ T-cells (*Fig.4J*).

Given the propensity of T_VM_ to produce IFN-γ during priming, and the role of IFN-γ in decreasing the avidity of the primary response leading to sub-optimal immunity, we investigated whether the presence of T_VM_ correlated with disease severity in humans. To do this, we made use of the COMBAT dataset, a blood atlas delineating innate and adaptive immune dysregulation in COVID-19 (Ahern et al. 2022). In humans, T_VM_ corresponds to a subset of CD8^+^ T effector memory RA (T_EMRA_) cells that expresses multiple killer Ig-like receptors (KIRs) and NKG2A/E (Thiele et al. 2020). We therefore focused on the T_EMRA_ clusters (*Fig.4K*) to identify T_VM_. Differentially expressed genes revealed that clusters 2, 3, 5 and 6 are characterized by multiple KIR and NKG2 expression (Dataset 1) and were therefore labelled as T_VM_ (*Fig.4L*). Analysis of the relationship between T_VM_ and disease severity revealed that enhanced frequency of T_VM_ is associated with severe COVID disease (*Fig.4M*). This is in agreement with our finding that CD8^+^ T-cells mainly sense IFN-γ produced by T_VM_, leading to decreased avidity of the primary response and a subsequently curtailed response during infection.

Altogether, our data demonstrate that paracrine IFN-γ sensing by CD8^+^ T-cell decreases the avidity of the primary response. Because T_VM_ is the subset producing IFN-γ during priming, we concluded that T_VM_ cells regulate the avidity of the CD8^+^ T-cell response.

### IFN-γ-sensing by CD8^+^ T-cells does not regulate CD8^+^ T-cell differentiation and TCR diversity

Our data so far demonstrate that IFN-γ sensing by CD8^+^ T-cells during priming lowers the avidity of the effector response, thereby limiting T-cell responses against intracellular pathogens. However, it was unclear whether IFN-γ intrinsically affected T-cell differentiation and fitness, or rather coordinated affinity, expansion, and differentiation. To address this question, we performed single-cell RNA- and TCR-sequencing on OVA-specific CD8^+^ T-cells from CD8-IFN-γR^KO^ and WT mice 9 days post-infection with LM-OVA. Using unsupervised hierarchical clustering, we identified 7 clusters (*Fig.S5A*), which we manually labelled based on known markers of effector and memory subsets (*Fig.S5B*). We distinguished 2 memory subsets based on *Cx3cr1* expression, *Cx3cr1*^*neg*^ being the least differentiated (Gerlach et al. 2016), one effector population and one highly cycling cluster (*Fig.5A-B*). WT and CD8-IFN-γR^KO^ clusters had similar gene expression profiles, showing that IFN-γ does not affect intrinsic effector or memory potential (*Fig.5C*). This agrees with the fact that WT and CD8-IFN-γR^KO^ T-cells exhibited similar cytotoxicity (*Fig1E-G*). We also observed that the IFN-γ signature of WT-cells at the peak of infection was low and not decreased by IFN-γR deletion (*Fig.S5C*), consistent with our finding that CD8^+^ T-cells sense IFN-γ during priming rather than during the effector stage.

**Figure 5:**
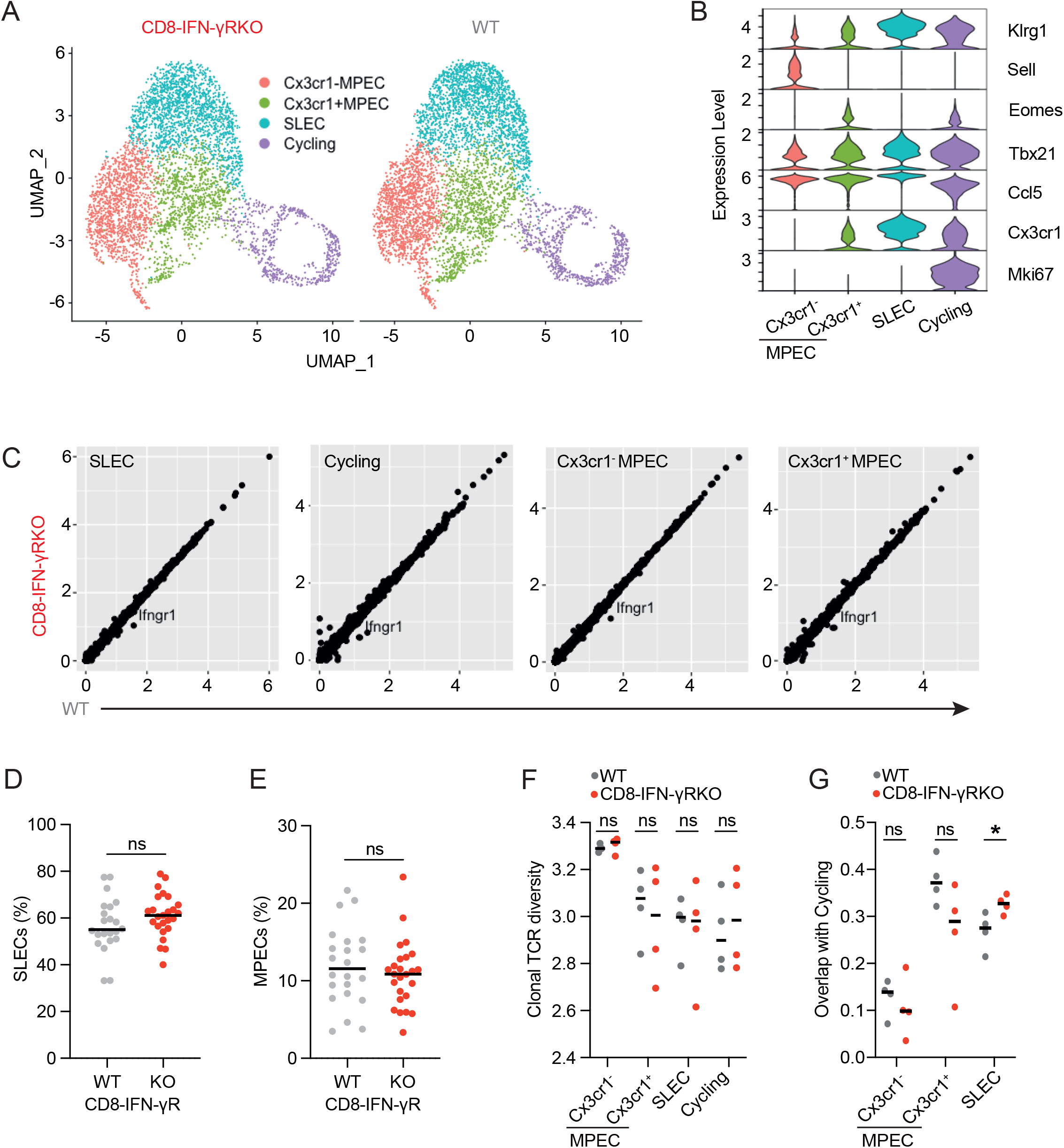
IFN-γR deletion in CD8^+^ T-cells does not regulate overall differentiation and TCR diversity. **(A-C, F-G)** CD8-IFN-γR^KO^ and WT mice were infected with LM-OVA, tetramer^+^ CD8^+^ T-cells were sorted from spleens 9 days post infection and subjected to scRNA-seq and scTCR-seq analysis (n=4). **A-** Graph-based clustering (n=5646 WT; n=4837 CD8-IFN-γR^KO^) of the assembled differentiation states. **B-** ViolinPlot shows the expression of selected markers. **C-** Plot shows the average gene expression of WT and CD8-IFN-γR^KO^ clusters. **(D-E)** CD8-IFN-γR^KO^ and WT mice were infected with LM-OVA and splenocytes were isolated 9 days post-infection to analyze the relative abundance of SLEC **(D)** and MPEC **(E)** cells by flow cytometry. **(F)** Relative TCR clonal diversity (Shannon index) by cell state and genotype extracted from scTCR-seq analysis. **(G)** Clonal overlap between the cycling cluster and the other clusters from scTCR-seq analysis. Data are pooled from 9 (**D-E**, n = 27) independent experiments. Error bars indicate the median. Comparison between groups was calculated using the two-tailed unpaired Student’s t-test. *P<0.05; **P<0.01; ***P<0.001; NS, not significant.

We then tested whether IFN-γ sensing would alter the differentiation of endogenous CD8^+^ T-cells. We did not detect any significant differences by flow cytometry (*Fig.5D-E*), where the proportion of SLECs (KLRGI^hi^CD127^lo^) (*Fig.5D*) and MPECs (KLRGI^lo^CD127^hi^) (*Fig.5E*) was identical between CD8-IFN-γR^KO^ and WT endogenous OVA-specific CD8^+^ T-cells at the peak of the response. It demonstrated that IFN-γ sensing does not regulate overall CD8^+^ T-cell differentiation. It was however worth noting that blocking overall IFN-γ during priming increased SLEC proportion (*Fig.S5D*) and conversely decreased MPEC proportion (*Fig.S5E*), which might be explained by the function of IFN-γ on other cell types, indirectly affecting CD8^+^ T-cell differentiation. For example, in models of vaccination, IFN-γ signaling in CD11b^+^ cells regulates CD62L expression on CD8^+^ T-cells (Sercan et al. 2010).

We then investigated whether the difference in avidity between WT and CD8-IFN-γR^KO^ T-cells was related to the use of a different TCR repertoire. Analysis of TCR usage showed that the OVA-specific T-cell repertoire following LM-OVA infection is unique for each mouse, regardless of the genotype (*Fig.S5F*), suggesting that TCR repertoires emerging after infection are qualitatively different, as observed during CMV infection (Schober et al. 2020). The increased avidity induced by IFN-γR deletion may be the result of the expansion and dominance of a few high-affinity T-cell clones, which would result in decreased TCR diversity. However, IFN-γ sensing by T-cells did not regulate TCR diversity (*Fig.5F*), indicating that differences in avidity are not solely due to an alternate priming of high-affinity T-cells. To get more insight into the function of IFN-γ on the relationship between clonal breadth and differentiation, we compared the TCR clonal overlap between the different subsets. The overlap between SLECs and the memory populations was limited, about 20%, and was not affected by IFN-γR deletion (*Fig.S5G*). Interestingly, analysis of the overlap between the cycling population and the other subsets revealed that inhibition of IFN-γ sensing enhanced the overlap between cycling and SLEC populations (*Fig.5G*), suggesting that the expansion of distinct clones was enhanced following IFN-γR deletion. Given our previous data demonstrating that high-affinity T-cells expanded more following IFN-γR deletion, we hypothesize that those clones were of high-affinity for their cognate peptide. This, however, was not enough to reduce TCR diversity, indicating that lower affinity T-cells were still efficiently primed and recruited. Similar results were obtained when we focused our analysis on the four most expanded TCRV_β_ chains (*Fig.S5H-I*).

We concluded that IFN-γ sensing does not affect T-cell differentiation *per se* and TCR diversity, but combined analysis of scRNA- and scTCR-seq suggests that IFN-γ sensing regulates the relationship between clonal expansion and T-cell subsets.

### IFN-γ-sensing by CD8^+^ T-cells increases the avidity of the memory response

Because low- and high-affinity T-cells are known to have distinct differentiation skewing (Knudson et al. 2013) and IFN-γ sensing by CD8^+^ T-cells lowers the avidity of the primary response, we hypothesized that IFN-γ may rather regulate the relationship between T-cell avidity and differentiation. To investigate this relationship, we first analyzed the differentiation state of adoptively transferred WT and IFN-γR^KO^ OT-I T-cells upon infection with LM expressing altered OVA peptides. IFN-γR deletion had limited effect on the differentiation of high-affinity T-cells, whereas it increased MPEC skewing of low-affinity T-cells (*Fig.6A-B*). This suggested that IFN-γR deletion enhanced memory formation of low-affinity T-cells. To test this, a small number of WT and IFN-γR^KO^ OT-I T-cells were admixed, and transferred in WT mice, which were challenged with LM expressing altered OVA peptides with varying affinities. After 60 days, mice were re-challenged with LM-OVA in order to analyze OTI expansion. Priming with high-affinity peptides led to a slight decreased frequency of IFN-γR^KO^ over WT OT-I T-cells during recall responses, as already suggested (Krummel et al. 2018). However, expansion of IFN-γR^KO^ OT-I T-cells primed with low-affinity peptides was increased compared to WT during recall responses (*Fig.6C*). This suggested that IFN-γ sensing by CD8^+^ T-cells increased the avidity of the memory response by limiting the inherent skewing in memory differentiation of low-affinity T-cells.

To test this hypothesis, we first analyzed the avidity of OVA-specific SLEC and MPEC CD8^+^ T-cells at the peak of the response. As expected, CD8-IFN-γR^KO^ SLECs were skewed towards higher tetramer binding compared with WT (*Fig.6D-E*). However, CD8-IFN-γR^KO^ MPECs displayed overall highly variable avidity at this time point (*Fig. 6D-F*). To specifically address whether IFN-γ affected the avidity of the memory response, LM-OVA infected CD8-IFN-γR^KO^ and WT mice were re-challenged after 60 days, and the expanded memory T-cell population was analyzed. CD8-IFN-γR^KO^ T-cells exhibited lower tetramer binding (*Fig.6G*), indicating that the memory compartment was skewed towards lower avidity in CD8-IFN-γR^KO^ mice. Similar results were observed following Influenza infection. We challenged WT and CD8-IFN-γR^KO^ mice with the Influenza strain X31-OVA. After 60 days, mice were re-challenged with another virus strain, PR8-OVA. This strategy was used because X31 and PR8 express the same NP_68_ T-cell epitope but different B-cell epitopes, ensuring the virus was not cleared by memory B-cells. Endogenous OVA-specific T-cells were indeed of lower avidity (*Fig.6H*) in the lung, but we could detect very few of those cells. We therefore also analyzed the avidity of the T-cell repertoire towards the dominant flu antigen NP_68_. Both in draining lymph nodes and in the lung, NP_68+_ CD8^+^ T-cells expanded less (*Fig.6I-J*) and were of lower avidity (*Fig.6K-L*) in CD8-IFN-γR^KO^ mice compared to WT mice. Altogether, we concluded that IFN-γ improves memory responses.

**Figure 6:**
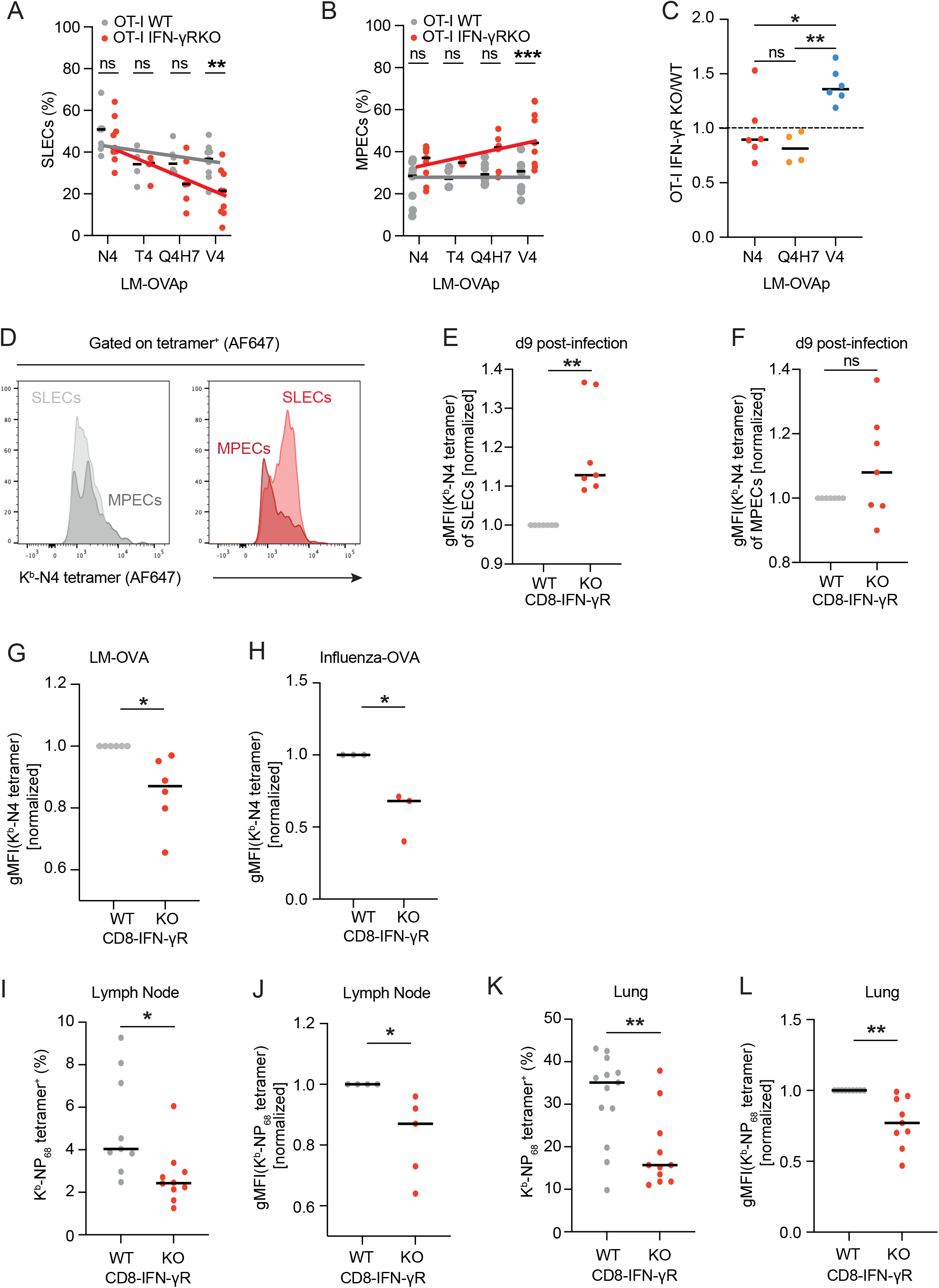
IFN-γR deletion in CD8^+^ T-cells lowers the avidity of CD8^+^ T-cell memory responses. **(A-C)** WT mice were transferred with 50,000 ad-mixed WT and IFN-γR^KO^ OT-I T-cells and infected with LM expressing the indicated OVA peptides. Spleens were isolated 9 days post-infection to analyze the relative abundance of SLEC **(A)** and MPEC **(B)** within the transferred OT-I population by flow cytometry. **C-** Mice were re-challenged after at least 60 days with LM-OVA and spleens were isolated 5 days after rechallenge to analyze OT-I T-cells expansion by flow cytometry. Data are expressed as the ratio between IFN-γR^KO^ and WT OT-I cell number. **(D-G)** CD8-IFN-γR^KO^ (red) and WT (grey) mice were infected with LM-OVA. **(D-E)** Splenocytes were isolated after 9 days and analyzed by flow cytometry. **D-** Representative histograms of tetramer staining within the SLEC and MPEC populations. **(E-F)** Normalized tetramer-staining of SLEC **(E)** and MPEC **(F). G-** Mice were re-challenged 60 days post-infection with LM-OVA and normalized tetramer gMFI was analyzed by flow cytometry after 5 days. **(H-L)** CD8-IFN-γR^KO^ (red) and WT (grey) mice were infected with X31-OVA. Mice were re-challenged 60 days post-infection with PR8-OVA. Tissues were harvested after 5 days and stained for CD8 T-cells and the indicated tetramer. **H-** Graph shows normalized N4-tetramer gMFI. **(I-J)** Graphs show the percentage of NP68 tetramer+ CD8+ T-cells in the LN (**I**) and lung (**J**). **(K-L)** Graphs show normalized NP_68_ tetramer gMFI in the LN **(K)** and lung **(L)**. Data are pooled from 3 (**A-C**, n = 6), 7 (**E-F**, each dot represents an experiment) or 5 (**G**, each dot represents an experiment) independent experiments. Tetramer-binding gMFIs of CD8-IFN-γRKO samples were normalized to mean WT expression within each independent experiment. Error bars indicate the median. Comparison between groups was calculated using the two-tailed unpaired Student’s t-test. *P<0.05; **P<0.01; ***P<0.001; NS, not significant.

Overall, our data demonstrate that IFN-γ sensing by CD8^+^ T-cells results in decreased avidity of the primary response, leading to sub-optimum control of infection. This is accompanied by increased avidity of the secondary response. T_VM_ are the main subset providing IFN-γ to other T-cells, linking innate signals to the regulation of CD8^+^ T-cell breadth, ensuring that the effector and memory pools are composed of a breadth of T-cell clones.

## Discussion

In this study, we described a critical role of IFN-γ as a paracrine modulator of CD8^+^ T-cell responses by coordinating T-cell expansion, differentiation, and avidity. Individual CD8^+^ T-cell clones exhibit a remarkable degree of functional and phenotypic plasticity, a prerequisite for effective adaptive responses (Kaech and Cui 2012; Buchholz, Schumacher, and Busch 2016; Rutishauser and Kaech 2010). However, such malleability necessitates regulatory mechanisms ensuring robustness at the population-wide level. Our data demonstrate that IFN-γ mediates such regulatory mechanisms, not only during infection but also during anti-tumor immunity (Mazet et al. 2023).

Regulating the avidity of the T-cell response is critical. Increased avidity in the same order of magnitude we observed for IFN-γR deletion has been correlated with resistance to viral infection in humans. Increased control of HIV-1 replication by CD8^+^ T-cells (Almeida et al. 2007) is correlated with increased avidity, and, consequently, HIV controllers display higher functional avidities of Gag-specific and HLA-B-restricted responses than non-controllers (Berger et al. 2011). Similar increased avidity has been observed in patients that cleared Hepatitis C compared to chronically infected patients (Yerly et al. 2008). This emphasizes the importance of understanding the mechanisms regulating the avidity of the T-cell response.

High-affinity T-cells have a competitive advantage over low-affinity T-cells, with ligand affinity determining the frequency of responding cells (Richard et al. 2018). Low-affinity T-cells are nevertheless efficiently recruited during an immune response (Zehn, Lee, and Bevan 2009), resulting in a physiological range of affinities of around 100-fold (Hebeisen et al. 2013; Dolton et al. 2018). Our data demonstrate that this is in part due to IFN-γ, improving low-avidity T-cell expansion and thereby lowering the threshold for low-affinity T-cells to enter the effector response, overcoming the selective advantage of high-affinity T-cells. Multiple hypotheses could explain the importance of recruiting lower-affinity T-cells during primary responses. During infection, too many high-avidity T-cells may drive immunopathology (Hillaire, Rimmelzwaan, and Kreijtz 2013). In addition, they become dysfunctional and exhausted in chronic conditions (Shakiba et al. 2021). Interestingly, in metastatic melanoma patients, PD-1^neg^ T-cell clones naturally present within an endogenous repertoire exhibit lower avidity TCRs than PD-1^+^ T-cell clones, exemplifying the tight relationship between T-cell avidity and exhaustion (Simon et al. 2016).

Lowering the avidity of the primary response may also be necessary to increase the avidity of the secondary response by allowing high-affinity T-cell clones, which are inherently biased to becoming effector, to also enter the memory pool, as shown here. While low-avidity T-cells are intrinsically skewed to become memory T-cells (Knudson et al. 2013) where they are important for memory responses towards mutated pathogens (van Gisbergen et al. 2011; Kavazović et al. 2020), IFN-γ ensures that high-avidity T-cells are also part of the memory pool, enabling efficient memory response towards both mutated and native pathogen.

Mechanistically, we found that IFN-γ is provided by T_VM_ to other CD8 T-cells during priming, serving as “Signal 3”. But how IFN-γ signaling coordinate the selection of high-versus low-affinity T-cells throughout the effector and memory pool is unclear. IFN-γ induces the expression of transcription factors known to regulate the threshold of T-cell activation, proliferation, and differentiation, such as c-Myc and T-bet (Gocher, Workman, and Vignali 2021; Richer, Nolz, and Harty 2013). Alternately, IFN-γ induces ICAM-1 expression, which has been shown to regulate memory formation (Gérard et al. 2013; Cox et al. 2013). Regulation of ICAM-1 expression in CD8^+^ T-cells could alter the nature of T-T communication and associated quorum sensing (Zenke et al. 2020). It is tempting to speculate that those factors may integrate with TCR signaling, but more studies are necessary to elucidate the mechanisms enabling this coordination.

Taken together, our work provides evidence that IFN-γ serve an important autoregulatory role by coordinating CD8^+^ T-cell avidity and differentiation, decreasing the avidity of the primary response while increasing the avidity of the memory response, ensuring a balanced control of short- and long-term immunity.

## Methods

### Mice

*CD8a*-Cre^GFP^ (JAX stock no.: 008766), IFN-γR^flox/flox^ (JAX stock no.: 025394), IFN-γ-GREAT-YFP (JAX stock no.: 017580), IFN-γ^KO^ (JAX stock no.: 002287), Nur77-GFP (JAX stock no.: 018974), *CD8a*KO (JAX stock no.: 002665) and OT-I (JAX stock no.: 003831) were purchased from Jackson Laboratory. C57BL/6J mice were purchased from Charles River, UK (JAX stock number: 000664). To generate CD8 IFN-γR^KO^ mice *CD8a*-Cre^GFP^ mice were crossed with IFN-γR^flox/flox^ mice, which were subsequently crossed to Rosa26-tdTomato mice (kind gift from the group of Tal Arnon). All experiments involving mice were conducted in agreement with the United Kingdom Animal (Scientific Procedures) Act of 1986 and performed in accordance to approved experimental procedures by the Home Office and the Local Ethics Reviews Committee (University of Oxford).

### Cell Isolation

Spleens were harvested from mice at indicated time points, mashed in 1x PBS and filtered through 70 μM filter. Splenocytes were resuspended in red lysis buffer (155 mM NH4Cl, 12 mM NaHCO3 and 0.1 mM EDTA in ddH_2_O) and incubated on ice for 5 minutes, before being washed in PBS twice.

For T-cell isolation, WT or OT-I CD8^+^ T-cells were isolated through negative separation from lymph nodes and spleens of 6- to 12-week-old WT or OT-I mice, using the MojoSort™ CD8^+^ T-cell isolation kit and magnets (Biolegend, #480008 and #480019). Isolated T-cells were resuspended in complete RPMI (RPMI 1640 [Gibco, #21870-076] supplemented with 2% FCS and 100x Penicillin-Streptomycin [Gibco, #10378-016]).

### Infection and Treatments

Mice were given an intravenous (i.v.) injection of 20*10^3^ colony-forming units (cfu) of LM expressing either a secreted form of OVA (LM-OVA) or one of the LM following strains expressing the indicated peptides (LM-N4, LM-T4, LM-Q4H7, LM-V4, LM-G4 or LM-gp33). LM strains were provided by Dietmar Zehn (TU Munich) expect LM-gp33, which was purchased from Nanjing Sungyee Biotech. Frozen down LM aliquots were expanded in Brain Heart Infusion (BHI) broth (Sigma, #53286-100G) and LM suspensions were plated on BHI agar plates (Sigma, #70138-500G). LM (200.000 cfu for 24h experiments; 20.000 cfu for d7-10 experiments) was injected intravenously when they were in exponential phase of growth. For memory responses, mice were rechallenged at least 60 days after the primary infection with 200.000 cfu.

In some experiments, OT-I T-cells (2-3×10^6^ cells or 50.000 cells for 24h or d7-10 experiments, respectively) were transferred into mice recipient by intravenous injection the days before infection.

In some experiments, mice received a single intraperitoneal (i.p.) injection 16-24 hours post infection of 75 μg of isotype matched control antibody (rat IgG1, BioXCell, #BE0088) or anti-Interferon-gamma (BioXCell, clone: XMG1.2). In some experiments 50 ug of isotype matched control antibody (rat IgG2a, BioXCell, #BE0089) or anti-NK1.1 (BioXCell, clone: PK136) was injected intraperitoneally two days and one day prior to infection, as well as on day 4 and 6 post-infection.

For subsequent staining of intracellular cytokines, mice were injected i.p. with 250 μg BFA 6 hours before being killed.

For primary infection with Influenza virus, mice weighing >20g were anaesthetized using isoflurane and intranasally administered with 4×10^4^ or 4×10^5^ PFU of X31-OVA influenza A virus in PBS. Mice were weighted for 14d following infection. For memory responses, mice were rechallenged at least 60 days after the primary infection with 10^6^ PR8-OVA.

### In vitro Activation and Treatment

For *in vitro* re-stimulation experiments, splenocytes were activated 7-10 days after infection with different concentrations of N4 peptide or phorbol 12-myristate 13-acetate (PMA) (2 ng/mL) and ionomycin (20 ng/mL). For *in vitro* stimulation experiments, naïve T-cells were seeded in 96-well U-bottom together with bone marrow-derived dendritic cells (BMDCs) loaded with the indicated OVA peptide (Proteogenix) at 10 ng/mL or antibodies. For experimental conditions including TCR stimulation, wells were pre-coated with anti-CD3ε (Biolegend, #145-2C11) prepared in PBS at 1 μg/mL for 2-3 hours at 37°C before being emptied for seeding. For ICAM-1 stimulation, wells were pre-coated with 5 μg/mL ICAM-1 (Biolegend, #553006) for 2-3 hours at 37°C before being emptied for seeding. Following coating with anti-CD3ε and/or ICAM-1, cytokine preparations were added to the wells according to the specified experimental conditions, which included different combinations of IL-12 (Biolegend, #577004), IL-18 (Biolegend, #767004), IL-15 (Biolegend, #566302), TNF (PeproTech, #315-01A-20uG), IL-33 (Biolegend, #580504), IFN-α (Biolegend, #752804), IFN-β (Biolegend, #581304), IL-2 (Biolegend, #575404), IL-1β (Biolegend, #575102), and IL-7 (PreproTech, #217-17). For co-stimulation, 1 μg/mL anti-CD28 (Biolegend, #37.51) was also added to wells pre-coated with anti-CD3ε. The cells were incubated overnight under the specified conditions at 37°C in 5% CO_2_ before being subjected to staining and flow cytometry analysis.

For subsequent staining of intracellular cytokines, cultured cells were treated with 7 μg/mL brefeldin A (BFA, ChemCruz, #sc-200861A) 30 minutes post-stimulation and incubated for 4.5 hours at 37°C in 5% CO2 before further processing.

### Generation of BM chimeras

IFN-γ^KO^ Recipient mice were irradiated with 4.25 Gy per cycle for two irradiation cycles, 4 hours apart. Bone marrow from IFN-γ^KO^, *CD8a*^KO^ or C57BL/6J mice were prepared from femur and tibia. Cell suspension was filtered through a 40 μM strainer shortly prior to intravenous injection of 100 μL (equivalent to 1.25 ×10^5^ cells/mouse). Mice were kept for two weeks post-transplantation on antibiotics (Enroflocaxin) in drinking water to avoid infection.

### In Vivo Cytotoxicity Assay

Isolated and washed splenocytes were resuspended in complete RPMI and divided into two 15 mL falcon tubes, each containing 1 mL splenocyte suspension. OVA peptide SIINFEKL (N4) was added to one splenocyte suspension to a final concentration of 10 μg/mL. Both the peptide-treated and untreated splenocytes were incubated at 37°C for 30 minutes and shaken half-way. The cells in both tubes were counted and resuspended to obtain 1 mL suspensions containing 10*10^6^ cells. Peptide-loaded and non-loaded cell suspensions were incubated with 1 μM eFluor 670 and 2 μM CFSE in 1x PBS, respectively. The dye-stained cell suspensions were centrifuged and resuspended in 1x PBS, before being mixed at a 1:1 ratio for injection into LM-OVA-infected recipient mice, as well as a naïve mouse used as control. Splenocytes were isolated from the spleens of infected and naïve mice 24h following injection, and cytotoxic killing by OVA-specific CD8^+^ T-cells was assessed through flow cytometry analysis. The degree of cytotoxicity is given as the percentage of targeT-cell lysis relative to the naive mouse, calculated by the following formula: 100-100*(R_infected_/Rn_aïve_), where the R values are equal to the % peptide-loaded population/% non-loaded population ratios in the infected and naive mice.

### Flow Cytometry

Single-cell suspensions obtained from spleen or cultured CD8^+^ T-cells were stained in V-bottom 96-well plates in flow cytometry buffer (2% FCS, 2 mM EDTA, and 0.02% sodium azide in 1x PBS). Live dead staining and surface staining was performed using Zombie NIR Fixable Viability Kit (Biolegend, #423106/423105), TruStain FcX^™^ (anti-mouse CD16/32, Biolegend, #101319) and fluorochrome-conjugated primary antibodies (Biolegend, Cell Signaling Technology or BD Biosciences), respectively. For experiments that included tetramer staining, cells were incubated with either Alexa Fluor 647- or BV421-conjugated, N4-specific MHC I tetramers (obtained from the National Institutes of Health Tetramer Core Facility [Emory University, Atlanta]) diluted 1:500 in flow cytometry buffer for 30 minutes at room temperature, prior to surface staining. If not otherwise indicated, all cells were fixed either in 4% paraformaldehyde (PFA) for 10 minutes at room temperature or using eBioscience^™^ Foxp3/Transcription Factor Staining Set (Invitrogen, #00-5523-00) for 30 minutes at 4°C. Intracellular transcription factor staining was performed after fixation and permeabilization using fluorescently labelled primary antibodies. Intracellular cytokine staining was performed after 15 minutes fixation using BD Cytofix/Cytoperm^™^ Fixation/Permeabilization Kit (BD Biosciences, #554714). Flow cytometry data were recorded on BD LSRII or FortessaX20 using DIVA software and analyzed using FlowJo^™^ software (Tree Star).

Surface markers used for flow cytometry analysis included: anti-CD8 (Biolegend, clone: 53-6.7), anti-CD4 (Biolegend, clone: RM4-5), anti-CD69 (Biolegend, clone: H1.2F3), anti-CD44 (Biolegend, clone: IM7), anti-CD49d (Biolegend, clone: R1-2), anti-NK1.1 (Biolegend, clone: PK137), anti-KLRGI (Biolegend, clone: 2F1/KLRF1). Antibodies used for intracellular cytokine staining included anti-IFN-gamma (Biolegend, clone: XMG1.2) and anti-TNF (Biolegend, clone: MP6-XT22).

### Cell binding avidity assays

For acoustic force spectroscopy of T cell avidity using the Z-Movi cell avidity analyser (LUMICKS, The Netherlands), microfluidic chips were functionalised with 1M NaOH followed coating with poly-L-lysine (Sigma Aldrich). Chips were kept dry in a 37°C incubator, with repeated aspiration of any residual liquid. For target cell monolayer seeding, microfluidic chips were rehydrated with prewarmed media. B16 tumour cells expressing OVA (70 – 80% confluency) were then seeded at a density of 100 million cells/mL and frequent checking under the microscope to ensure no bubble formation and appropriate seeding density. Cells were incubated for 3h at 37°C, with fresh change of media in between. Experiments were performed with sorted tetramer^pos^ cells isolated from 4 WT and 4 CD8-IFN-γR^KO^ mice 9 days after LM-OVA infection on each chip. Sorted cells were cultured for 3 days prior to measuring cell avidity. Effector T cells were stained with CellTrace Far Red Proliferation Kit (ThermoFisher Scientific) according to manufacturer’s instructions. The labelled cells were then seeded at a density of 10 million cells/mL onto the microfluidic chip containing the target cell monolayer and incubated for 15 min prior to increasing force application to measure cellular avidity. The 2 different types of effector cells were evaluated on the same microfluid chip, and the order was randomised between chips on repeated runs. Image-based automated detection of T cell detachment was performed using the Oceon software (LUMICKS) and analysis was conducted according to manufacturer recommendations.

### Single-cell RNA sequencing

For CD8^+^ T-cell sequencing during priming: CD8^+^ T-cells from 3 naïve mice and 3 mice infected with LM-OVA for 24h were sorted from splenocytes, cryopreserved in 20% FBS and 10% DMSO in RPMI and further processed by Single Cell Discoveries, Netherlands. Samples were further processed in accordance with 10x Genomics single cell protocols. Single cell libraries were prepared using the Chromium 3 ′ v2 platform (10× Genomics, Pleasanton, CA) following the manufacturer’s protocol. In brief, single cells were encapsulated into gel beads in emulsions (GEMs) in the GemCode instrument followed by cell lysis and barcoded reverse transcription of RNA, amplification, shearing and 3′ adaptor and sample index attachment. Approximately, 5000 to 7000 cells were recovered. Libraries were sequenced on the Illumina NovaSeq 6000. Read mapping, alignment to GRCm38 and quantitation of sample count matrices was performed with the 10x Genomics Cell Ranger pipeline (v 4.0.0).

For CD8^+^ T-cell sequencing at the peak of the primary response: tetramer^pos^ CD8^+^ T-cells from 4 WT and 4 CD8 IFN-γR^KO^ mice infected with LM-OVA for 9 days were sorted from splenocytes. Each mouse from each genotype was labelled with TotalSeq^™^ Hashtags (Biolegend) and mixed. Approximately 20,000 cells per sample were loaded onto the 10X Genomics Chromium Controller (Chip K). Gene expression, feature barcoding and TCR sequencing libraries were prepared using the 10x Genomics Single Cell 5’ Reagent Kits v2 (Dual Index) following manufacturer user guide (CG000330 Rev B). The final libraries were diluted to ∼ 10nM for storage. The 10nM library was denatured and further diluted prior to loading on the NovaSeq6000 sequencing platform (Illumina, v1.5 chemistry, 28bp/98bp paired end for gene expression and feature barcoding, 150bp paired end for TCR libraries).

### scRNA-sequencing analysis

Datasets were analyzed using Seurat version 4.0.5 (Hao et al., 2021). For CD8^+^ T-cell sequencing during priming, we filtered out T-cells having less than 600 and more than 5,000 detected genes, cells in which mitochondrial protein-coding genes represented more than 10% of UMI. Cells were then further filtered based on the expression of *Cd2, Cd8a* and *Cd8b1*. Samples were then integrated with the IntegrateData function and normalized with the scTransform function of Seurat and variation associated with mitochondrial and ribosomal UMI percentage were regressed out. Principal components were calculated using the top 3,000 variable features. These genes were used as input for principal component analysis (PCA), and significant PCs (n = 30) identified using Seurat (“JackStraw” test and “Elbowplot”). Clustering was performed with the Louvain algorithm (n = 30 PCs, resolution = 0.3).

For CD8^+^ T-cell sequencing at the peak of the primary response, we filtered out T-cells having less than 500 and more than 5,500 detected genes, cells in which mitochondrial protein-coding genes represented more than 5% of UMI and cells in which the percentage of largest genes was more than 15% of UMI. Cells were then further filtered based on the expression of *Cd2, Cd8a* and *Cd8b1*. Samples were then integrated with the IntegrateData function and normalized with the scTransform function of Seurat and variation associated with mitochondrial UMI percentage were regressed out. Principal components were calculated using the top 3,000 variable features. These genes were used as input for principal component analysis (PCA), and significant PCs (n = 20) identified using Seurat (“JackStraw” test and “Elbowplot”). Clustering was performed with the Louvain algorithm (n = 20 PCs, resolution = 0.3). A large cluster corresponding to a contamination with naïve CD8 T cells was removed and dataset was re-normalized, scaled and PCA and UMAP were re-calculated.

For differential expression analysis, NormalizeData and ScaleData were run on the RNA assay of the integrated data. Significant differentially expressed genes between clusters were identified using the “FindAllMarkers” function, Wilcoxon test and selecting markers expressed in at least 25% of cells. Significant differentially expressed genes between stimulation within clusters were identified using the “FindMarkers” function, Wilcoxon test and selecting markers expressed in at least 25% of cells. Pathway analysis was performed with Fast gene set enrichment analysis (fgsea), using the Gene Ontology or the Reactome pathway repositories. IFNg signature (Ifngr2, Stat1, Stat2, Irf1, Irf2, Irf7, Cxcl9, Cxcl10, Cxcl11) was computed using the package UCell (Andreatta and Carmona 2021).

### For TCR sequencing analysis

paired chain TCR sequences were obtained through targeted amplification of full-length V(D)J segments during library preparation. Sequence assembly and clonotype calling was done through cellranger’s immune profiling pipeline (cellranger multi). TCR profiling on filtered contig annotations was done using R package scRepertoire version 1.1.4 (Borcherding, Bormann, and Kraus 2020). Only cells for which both TCRa and TCRb could be identified were used. Clone calling was done for each sample set independently before integration in the Seurat object. Diversity was done by calculating the Shannon Index for each mouse, using the function clonalDiversity in scRepertoire.

### Statistical Analysis

One-way ANOVA was selected for pairwise comparisons across multiple experimental conditions and unpaired student’s t-tests were used to compare two conditions for statistical significance. EC_50_ between groups was calculated by fitting a 3-parameter fixed-slope Hill function and confirming good fit by the R square function and visual inspection before performing a F-test to compare the model parameters. Data were considered statistically significant when p < 0.05. Data are presented as mean ± SEM or ± SD. Statistical analysis was performed using GraphPad Prism 7 software. Ns=non-significant, *P < 0.05, **P < 0.001, ***P < 0.0002 and ****P < 0.0001.

## Data availability

The mouse scRNAseq and scTCRseq data generated in this study will be deposited in the GEO database. Datasets reused in this study: EGAS00001005507. All data are included in the Supplemental Information or available from the authors upon reasonable requests, as are unique reagents used in this Article.

## Acknowledgments

We thank Tal Arnon, Michael Dustin, Vivian W. C. Lau, Anne Chauveau and Mariana Borsa for critical reading of the manuscript. We thank the Lumicks team, in particular Brittany Wingham, for assistance with cell binding avidity experiments and JonathanWebber for assistance with cell sorting. We thank the NIH Tetramer facility for all the tetramers used in this study. This work was supported by the BBSRC (BB/R015651/1 to A.G), Cancer Research UK (CR-UK) (C5255/A18085 through the Cancer Research UK Oxford Centre and 29549 to A.G); the Kennedy Trust for Rheumatology Research (KENN151607 and KENN202112 to A.G), John Fell Funds (0006162 to A.G), MLSTF funds and Kennedy Trust Price Studentship (to L.F.K.U).

