## Supplementary figures and images for "Integration of Avidity and Differentiation is enabled by CD8^+^ T-cell sensing of IFN-γ"

### Suppl Figures

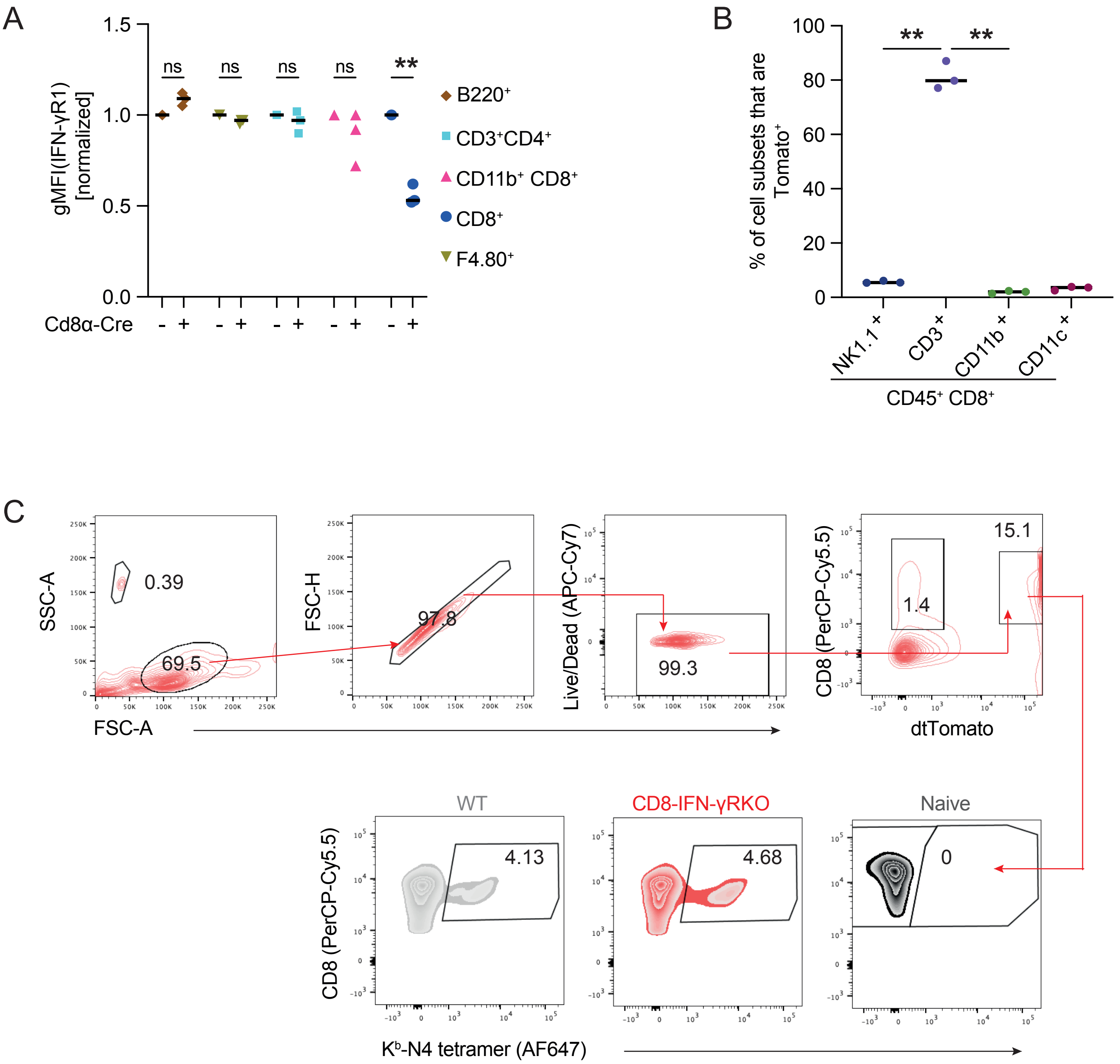

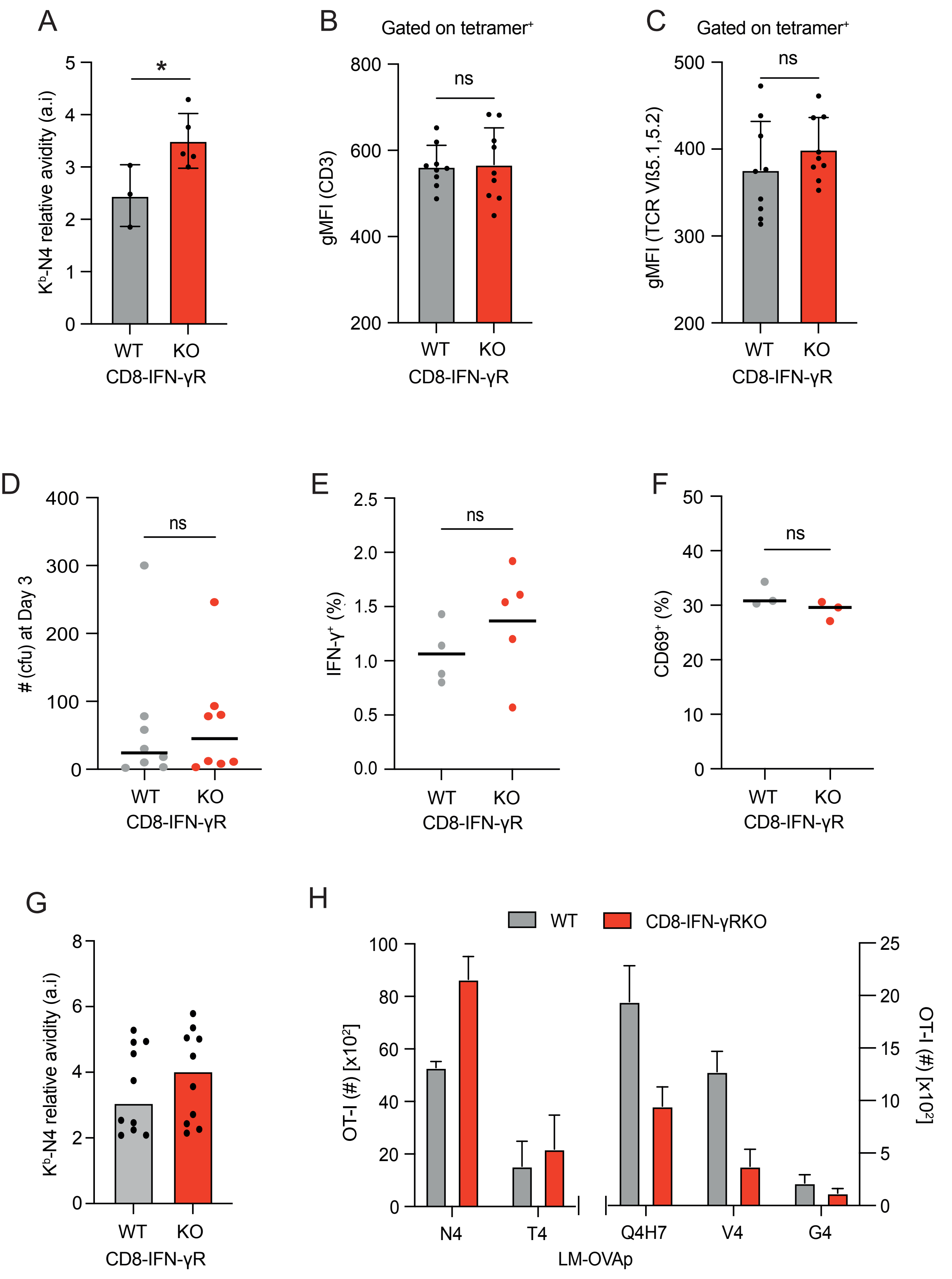

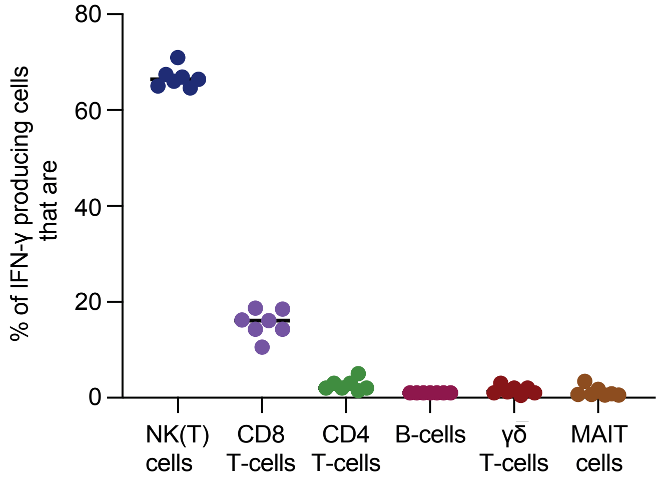

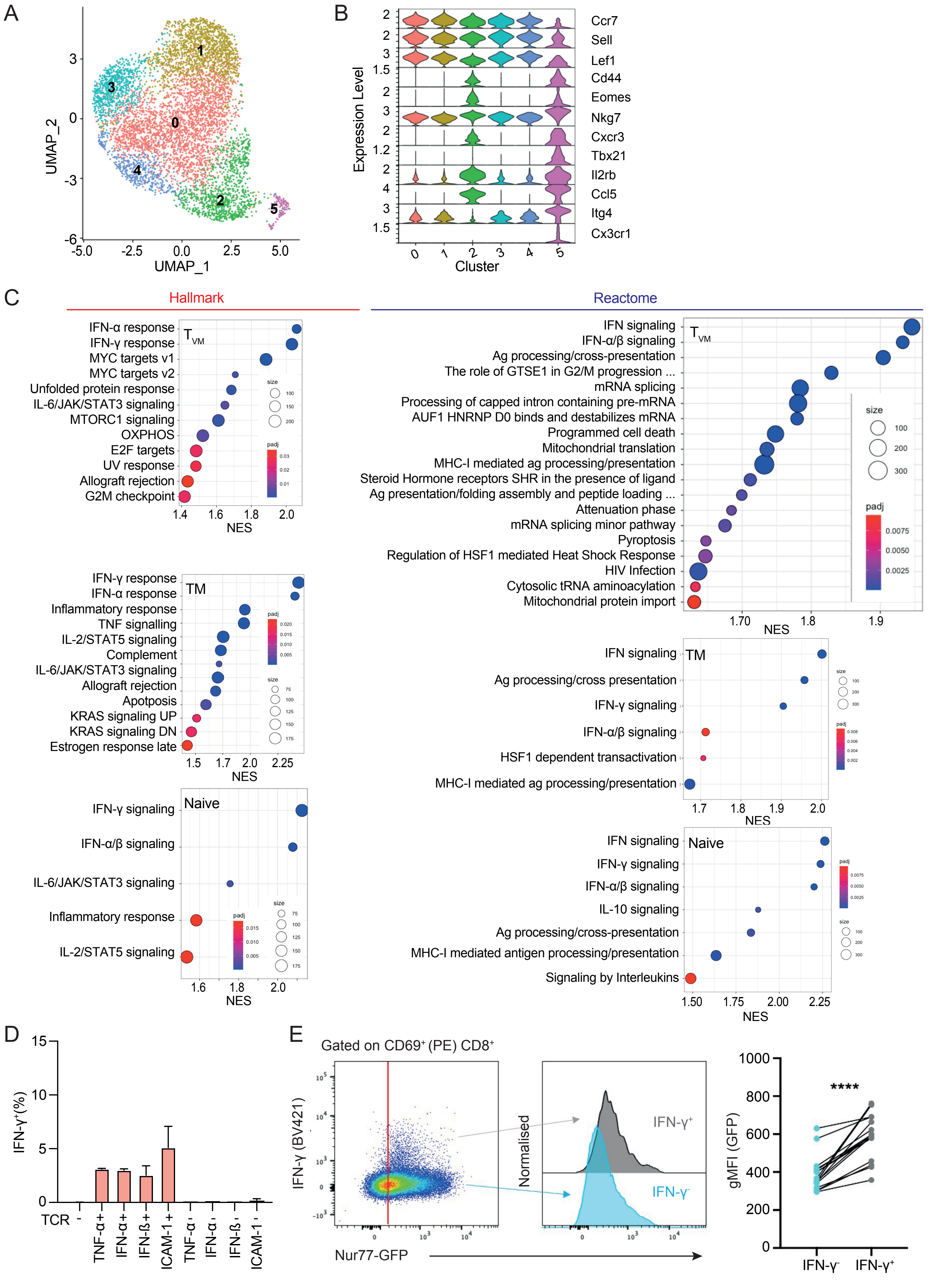

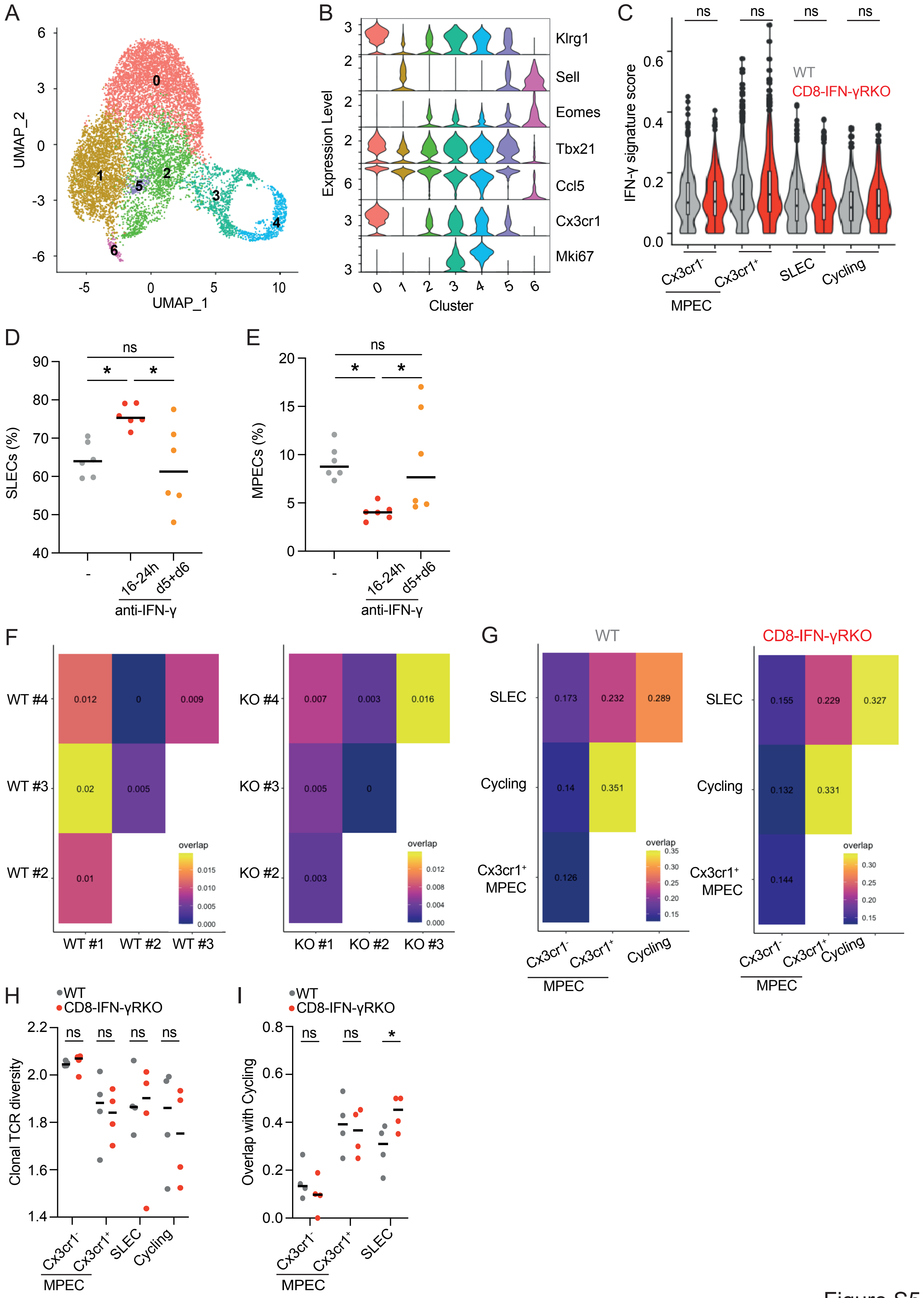
