## Supplementary material for "Integration of Avidity and Differentiation is enabled by CD8^+^ T-cell sensing of IFN-γ": Suppl Figure legends

### Supplementary Figure legends

#### Figure S1: Specific deletion of IFN-γR in CD8^+^ T-cells

**(A-B)** Cell-specific deletion of IFN-γR in different immune cell subtypes of CD8-IFN-γR^KO^ or WT mice was assessed either by flow cytometry analysis of IFN-γR1 **(A)** or tdTomato expression **(B)**. **C-** Representative dot plots of gating strategy for endogenous OVA-specific, tetramer^+^ CD8^+^ T-cells. Data are representative of 2 (**A-B,** n = 3) independent experiments. Error bars indicate the median **(A-B)**. Comparison between groups was calculated using the two-tailed unpaired Student’s t-test **(A-B)**. *P<0.05; **P<0.01; ***P<0.001; NS, not significant.

#### Figure S2: IFN-γR deletion in CD8^+^ T-cells increases the avidity of the primary response.

**(A-G)** CD8-IFN-γR^KO^ (red) and WT (grey) mice were infected with LM-OVA. **(A-C)** Spleens were isolated and tetramer^+^ CD8^+^ T-cells were analyzed after 9 days. **A-** Relative avidity of tetramer^+^ CD8^+^ T cells, calculated by dividing tetramer MFI by CD3 MFI **(B)**. **C-** Quantification of TCR V_β_5.1,5.2 expression of tetramer^+^ CD8^+^ T-cells. **D-** Mice were isolated after 3 days and bacterial load was analyzed. **E-** Mice were treated with BFA 6hrs before harvest. Spleens were isolated after 24hrs and IFN-γ expression by splenocytes was analyzed by flow cytometry. **F-** Spleens were isolated after 24hrs and CD69 expression on CD8^+^ T-cells was analyzed by flow cytometry. **G-** Spleens were isolated and tetramer^+^ CD8^+^ T-cells were analyzed after 5 days. Graph shows the relative avidity of tetramer^+^ CD8^+^ T-cells. **H-** WT mice were transferred with 50,000 ad-mixed WT (grey) and IFN-γR^KO^ (red) OT-I T-cells and infected with LM expressing the indicated OVA peptides. Spleens were isolated 9 days post-infection. Graph shows the absolute OT-I numbers, quantified by flow cytometry. Data are representative of 3 (D-F, n = 3-6) or pooled from 5 (A-B, each dot represents an experiment), 3 (C-D, G, n = 6-9) independent experiments. Error bars indicate the median (D-F) or mean ± s.e.m (D-H). Comparison between groups was calculated using a Two-way ANOVA and Šidák’s multiple comparison test. *P<0.05; **P<0.01; ***P<0.001; NS, not significant.

#### Figure S3: Immune cells producing IFN-γ during priming.

GREAT mice were infected with LM-OVA. Spleens were isolated after 24 hours. Splenocytes were stained for CD19, CD3, CD8, CD4, NK1.1, MR1, gd TCR. Immune populations among IFN-γ (YFP)^+^ splenocytes were analyzed by flow cytometry.

#### Figure S4: IFN-γ-sensing by CD8^+^ T-cells is paracrine and enabled by T_VM_

**(A-B)** CD8^+^ T-cells from naïve mice or infected with LM-OVA for 24h were sorted and subjected to scRNA-seq analysis (n=3). **A-** Graph-based clustering (n=3224 ctrl; n=3495 LM-OVA) of the identified clusters. **B-** ViolinPlot shows the expression of selected markers. **C-** Pathway analysis between control and LM-OVA infected samples for TM (top panel), T_VM_ (middle panel) or naïve (bottom panel) CD8^+^ T cells. Graphs show the Normalized Enrichment Score (NES) for the Hallmark pathways (left panels) and Reactome pathways (right panels). **D-** CD8^+^ T cells were stimulated in vitro with TCR/CD28 antibodies, IL-12, IL-18, ICAM-1, TNF, IL-15 or Type I IFN as indicated. IFN-γ expression was analyzed by flow cytometry after 24h. **E-** Nur77-GFP mice were infected with LM-OVA and Nur77-GFP and IFN-γ expression was evaluated by flow cytometry 24h post-infection. Shown are a representative flow plot (left panel) and histograms (middle panel) of Nur77-GFP and IFN-γ expression and the quantification (right panel) of Nur77-GFP expression in IFN-γ^neg^ and IFN-γ^pos^ CD8^+^ T cells. Data are pooled from 3 (**D-E,** n = 6-9) independent experiments. Error bars indicate the mean ± s.e.m **(D)**. Comparison between groups was calculated using a Two-way ANOVA and Šidák’s multiple comparison test. *P<0.05; **P<0.01; ***P<0.001; NS, not significant.

#### Figure S5: IFN-γ-sensing by CD8^+^ T-cells is paracrine and enabled by T_VM_

**(A-C,F-I)** CD8-IFN-γR^KO^ and WT mice were infected with LM-OVA, tetramer^+^ CD8^+^ T-cells were sorted from spleens 9 days post infection and subjected to scRNA-seq and scTCR-seq analysis (n = 4). **A-** Graph-based clustering (n = 5646 WT; n = 4837 CD8-IFN-γR^KO^) of initial clusters**. B-** ViolinPlot shows the expression of selected markers. **C-** Analysis of IFN-γ signalling signature of CD8-IFN-γR^KO^ (red) and WT (gray). Graph shows signature scoring by cell type and genotype. **(D-E)** WT mice were infected with LM-OVA and mice were either left untreated (grey) or treated with anti-IFN-γ 16-24 hours (red) or at day 5 and 6 (orange) post-infection. Splenocytes were isolated 9 days post infection and cell subsets were analyzed by flow cytometry using the markers KLRG1 and CD127. Graphs show relative the proportion of SLECs (KLRG1^+^ CD127^-^) **(D)**, and MPECs (KLRG1^-^ CD127^+^) **(E)** of tetramer^+^ CD8^+^ T-cells. **F-** Graph shows the frequency of TCR overlap between mice from combined scRNA- and scTCR-seq. **G-** Graph shows the frequency of TCR overlap between clusters from combined scRNA- and scTCR-seq. (H-I) Data was subsetted on the four most represented TCR Vb chains. **H-** Relative TCR clonal diversity (Shannon index) by cell state and genotype extracted from scTCR-seq analysis. **I-** Clonal overlap between the cycling cluster and the other clusters from scTCR-seq analysis. Data are pooled from 3 (**D-E,** n = 6-12) independent experiments. Error bars indicate the median. Comparison between groups was calculated using the two-tailed unpaired Student’s t-test **(D-E)**. *P<0.05; **P<0.01; ***P<0.001; NS, not significant.

### Supplementary Dataset 1

Differentially expressed genes between the different TEMRA clusters from the COMBAT dataset.
